# Conformational Barrier-Driven Flux Redistribution across Competing Catalytic Pathways Unifies Diverse Enzyme Kinetic Regimes

**DOI:** 10.64898/2026.09.21.753117

**Authors:** Rohon Mitra, Biman Jana

## Abstract

Enzymes are central to biological function, catalyzing the chemical transformations that sustain metabolism, signalling, molecular transport, and cellular regulation. Understanding their catalytic activity requires connecting molecular structure and conformational dynamics to measurable reaction kinetics. Classical Michaelis–Menten kinetics describes turnover through a dominant catalytic pathway approaching a single saturation limit, whereas conformational-selection, induced-fit, allosteric, and dynamic-disorder models account for additional complexity arising from conformational exchange and heterogeneous catalytic states. Experimental studies further reveal concentration-dependent changes in pathway usage, cooperative or sigmoidal responses, transitions between distinct turnover regimes, and intermittent single-molecule activity. These behaviours are typically treated within separate kinetic descriptions, leaving their mechanistic relationship within a common physical picture incompletely resolved. Here, we formulate a minimal conformational free-energy landscape with interconverting enzyme states and competing catalytic routes to examine whether these apparently distinct kinetic signatures can arise from a shared physical mechanism. In this framework, substrate availability and conformational barriers redistribute catalytic flux between slower and faster pathways, thereby altering both pathway occupancy and turnover times. Stochastic Gillespie simulations together with deterministic mean-first-passage-time analysis show that different parameter regimes generate low- and high-turnover states, concentration-dependent kinetic crossovers, burst–halt intermittency consistent with single-molecule observations, and effectively Michaelis–Menten-like or sigmoidal allosteric-like responses. The same underlying landscape can therefore produce qualitatively different kinetic behaviours depending on how conformational exchange and substrate capture partition catalytic flux. It also explains how finite substrate windows can mask underlying bimodal behaviour. These results provide a physically interpretable framework linking conformational dynamics, pathway selection, and experimentally observed enzyme kinetics.

## Introduction

Enzymes govern the chemical transformations that sustain biological function, enabling reactions to proceed with remarkable efficiency and selectivity under physiological conditions. [1–4] Through their participation in metabolism, energy conversion, signalling, molecular transport, and biomolecular synthesis and degradation, enzymatic activity is integrated into nearly every aspect of cellular regulation. Understanding how enzymes achieve and regulate this catalytic activity is therefore central to establishing the physical principles that connect molecular structure, conformational dynamics, and biological function. [5–8]

Enzyme kinetics provides a quantitative description of how catalytic activity changes with substrate availability and other reaction conditions. By relating reaction velocity to experimentally controlled variables, kinetic measurements can reveal characteristic turnover times, saturation behavior, rate-limiting steps, and changes in the population of catalytically relevant states. [5, 9] Such measurements therefore provide an important bridge between the macroscopic rate of an enzymatic reaction and the microscopic processes of substrate binding, conformational rearrangement, chemical conversion, and product release that collectively determine catalytic turnover. [7, 10–12] Several kinetic, biophysical, and statistical-mechanical frameworks have been developed to describe how molecular events within an enzyme are translated into measurable catalytic rates. These approaches differ in the level at which the catalytic process is represented, ranging from effective rate laws to models that explicitly account for free-energy barriers, conformational transitions, and redistribution among kinetically distinct states. [13–16] Within this broader theoretical landscape, Michaelis–Menten kinetics provides the classical reference description of enzyme turnover, in which substrate binding leads to formation of an enzyme–substrate complex followed by catalytic conversion and product release. [17–19] The resulting rate–substrate relation is hyperbolic and approaches a single saturation limit at high substrate concentration, describing the characteristic response expected when turnover is governed predominantly by one effective catalytic sequence.

Beyond this classical picture, several kinetic, biophysical, and statistical-mechanical frameworks have been developed to account for additional levels of enzymatic complexity. Transition-state theory and Kramers-type barrier-crossing descriptions relate catalytic rates to the free-energy barriers separating reactant, intermediate, transition, and product states, thereby linking turnover to molecular interactions, solvent reorganization, and conformational fluctuations. [8, 13, 15, 20, 21] At the level of substrate recognition, conformational-selection and induced-fit mechanisms describe two limiting routes by which the catalytically competent complex can be formed. In conformational selection, the enzyme samples a pre-existing ensemble of states and the substrate preferentially binds a competent conformation, whereas in induced fit, substrate association precedes the structural rearrangement required to reach the active state. [22–25] These mechanisms are not necessarily mutually exclusive, and their relative contribution can depend on the kinetic flux through the available binding pathways. [26–29] Conformational redistribution also forms the basis of cooperative and allosteric kinetic descriptions. The Monod–Wyman–Changeux model describes concerted transitions between conformational states with different ligand affinities, whereas the Koshland–Némethy–Filmer model considers sequential ligand-induced conformational changes among interacting subunits. [30–33] Such coupling can give rise to sigmoidal rate–substrate responses, in contrast to the hyperbolic saturation characteristic of the Michaelis–Menten limit. Related cooperative behavior can also arise in monomeric enzymes through slow interconversion between conformational states with distinct binding or catalytic properties. [34–40]

Experimental studies have further revealed concentration-dependent kinetic behavior that extends beyond the responses described by conventional kinetic models. This behavior is particularly evident in chaperonin systems, where ligand concentration can shift the population among kinetically distinct conformational states. In GroEL, ATP-dependent redistribution among the structurally unexpanded, hydrophobic substrate-binding state (TT), the asymmetric, GroES-capped folding state (TR), and the transient, fully expanded dual-ring state (RR) gives rise to two distinct transitions in ATPase activity, whereas the eukaryotic chaperonin containing tailless complex polypeptide 1 (CCT) exhibits separate kinetic phases associated with different conformational transitions. [41–45] Concentration-dependent changes can also extend beyond state redistribution to the relative use of alternative reaction pathways. In coupled binding–conformational systems such as dihydrofolate reductase (DHFR), for example, ligand binding and catalytic progression involve redistribution among conformational substates and kinetically distinct routes as reaction conditions change. [46–49] Together, these observations demonstrate that ligand concentration regulates not only the overall substrate occupancy but also the relative population of competing conformational pathways. This redistribution among pathways leads to two distinct saturation regimes, characterized by a lower kinetic rate under substrate-limited conditions and an enhanced kinetic rate at high substrate availability. A complementary form of nonclassical behavior emerges at the single-molecule level, where catalytic activity can vary substantially with time even under fixed reaction conditions. Measurements on individual cholesterol oxidase molecules have revealed slow fluctuations in turnover activity together with correlations between successive catalytic events, indicating conformational memory that is largely averaged out in ensemble measurements. [50–52] In single-molecule studies of (*β*)-galactosidase at high substrate concentration, turnover events occur in pronounced clusters separated by extended intervals of low activity. [53, 54] This temporal intermittency gives rise to a characteristic burst–halt pattern, consistent with stochastic transitions among conformational states possessing different catalytic rates. [55–57] These findings reveal a complementary form of nonclassical kinetics, in which temporal transitions among conformational states with distinct catalytic activities occur in addition to the concentration-dependent redistribution of catalytic flux described above.

In practice, these nonclassical kinetic responses are often characterized through fitting and statistical analysis of the measured behavior. [58, 59] Complex kinetic responses are typically characterized by fitting the experimental data to multistep reaction schemes or combinations of exponential components, [58] while single-molecule dynamics are quantified through waiting-time distributions, [60, 61] turnover-time statistics, and temporal correlations between successive catalytic events. [62, 63] Such approaches are effective in identifying distinct kinetic phases, characteristic timescales, and changes in apparent catalytic activity, but they usually describe the particular kinetic regime being probed rather than providing a common mechanistic origin for the full response. [64] As a result, classical saturation, cooperative or sigmoidal behavior, concentration-dependent switching between catalytic regimes, and intermittent single-molecule activity are generally treated within separate descriptions. [65] A broader mechanistic framework that connects these responses within the same conformational landscape therefore remains limited. In particular, such a framework should account for how substrate concentration, conformational exchange, and barrier crossing redistribute catalytic flux between competing pathways and thereby generate different kinetic regimes, both in the steady-state response and in the temporal organization of individual turnover events.

To unite these diverse kinetic manifestations within a single physical framework, we formulate a minimal conformational free-energy landscape for enzyme turnover. Within this landscape, the enzyme interconverts between distinct relaxed and tensed conformational states that are separated by conformational barriers and connect differently to substrate-bound intermediates, thereby defining competing routes for substrate entry and catalytic turnover. The present analysis shows that increasing substrate concentration shifts catalytic flux from the slower to the faster conformational pathway, giving rise to distinct low- and high-turnover regimes connected by a kinetic crossover. The resulting description relates changes in the conformational landscape to redistribution of catalytic flux between competing pathways and to variations in the residence time of slower catalytic states, thereby shaping the observed kinetic response. As the kinetic separation between the competing pathways increases, this redistribution is expressed not only in the steady-state response but also in the temporal organization of turnover, where stochastic trajectories exhibit pronounced burst–halt intermittency. Moreover, under limiting conditions, the same conformational landscape can recover effectively single-saturating Michaelis–Menten-like or sigmoidal allosteric-like responses over finite substrate ranges, indicating that these apparently distinct kinetic behaviors can emerge from different operating regimes of the same underlying catalytic network.

In a broader biophysical context, catalytic regulation can be viewed as a redistribution of population and flux over an underlying conformational free-energy landscape, rather than as a consequence of active-site chemistry alone. [66, 67] Perturbations that reshape this landscape can alter the accessibility of functional states and redirect the pathways through which catalysis proceeds. [68,69] The present description therefore connects enzyme kinetics to the broader ensemble-based picture of allostery, in which changes in conformational populations and exchange are translated into distinct functional outputs. [70, 71]

## Model

We describe enzyme turnover using a free-energy landscape framework that connects the classical Michaelis–Menten (MM) scheme with an extended non-MM description. In the MM landscape, shown in Fig. 1A, catalysis follows one dominant route: substrate capture is followed by barrier crossing, product formation, and product release. This gives a single saturation limit at high substrate concentration, corresponding to one effective turnover rate. The extended landscape in Fig. 1B represents enzymes whose turnover cannot be described by a single MM-like route. In this description, the enzyme can interconvert between two conformational states, denoted by *E*_*R*_ and *E*_*T*_ , and substrate can enter the catalytic cycle through more than one route. Consequently, the observed rate is determined not only by substrate binding, but also by how catalytic flux is distributed between the available pathways. As substrate concentration changes, this flux distribution can shift, producing a transition from a lower saturation regime to a higher saturation regime in the rate-substrate response. The model is therefore constructed to describe substrate-dependent switching between distinct kinetic regimes. For this extended non-MM landscape, the elementary kinetic reactions, rate constants, and model parameters are summarized below.

**Figure 1.**
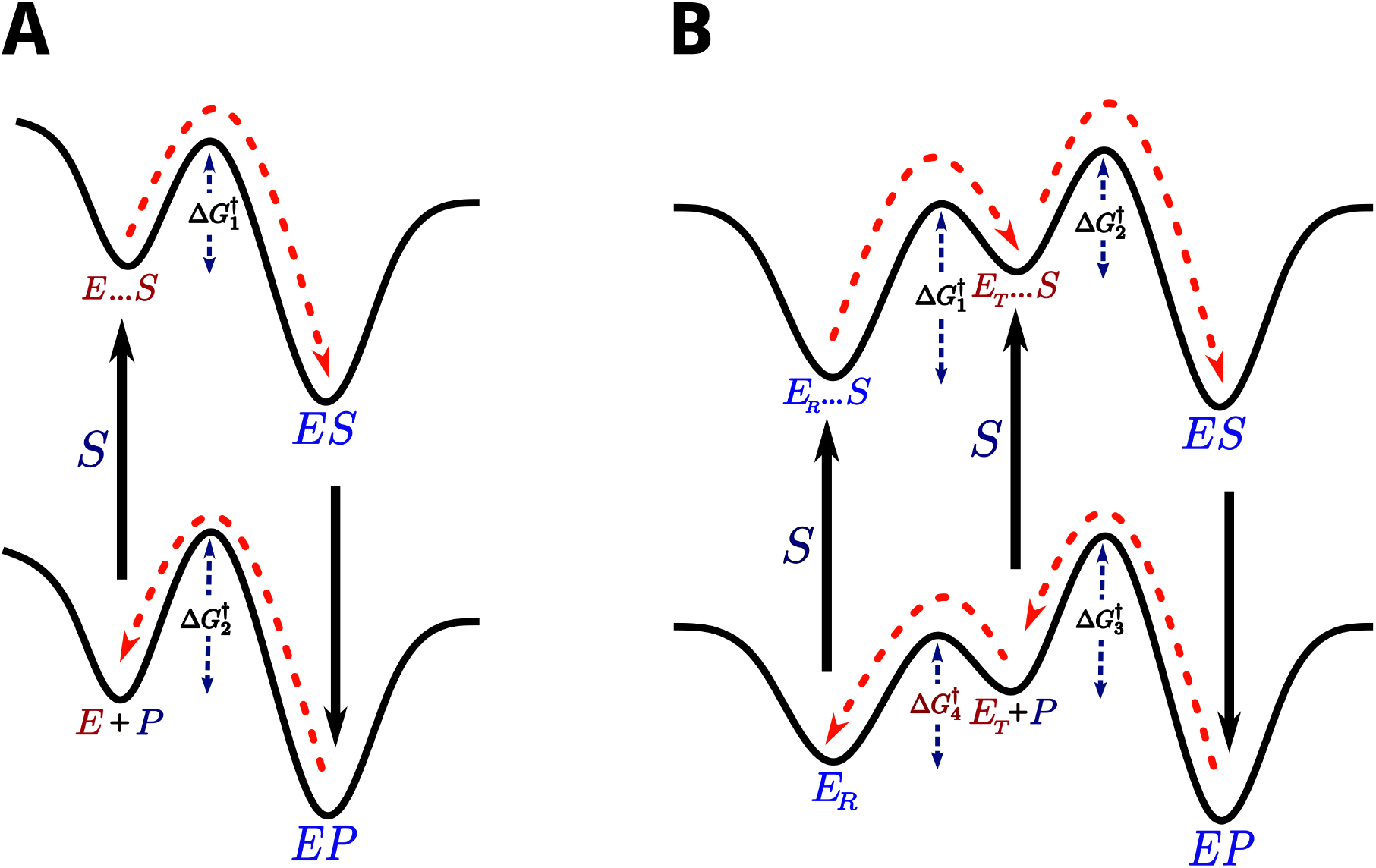
Schematic free-energy landscapes for single-pathway and conformationally coupled enzyme turnover. **(A)** Classical Michaelis–Menten kinetics, in which substrate binding, chemical conversion, and product release proceed through one dominant catalytic sequence. **(B)** Extended landscape with relaxed and tensed conformations, parallel substrate-entry routes, and separate activation barriers governing conformational exchange, catalytic progression, and pathway selection.

**Figure 2.**
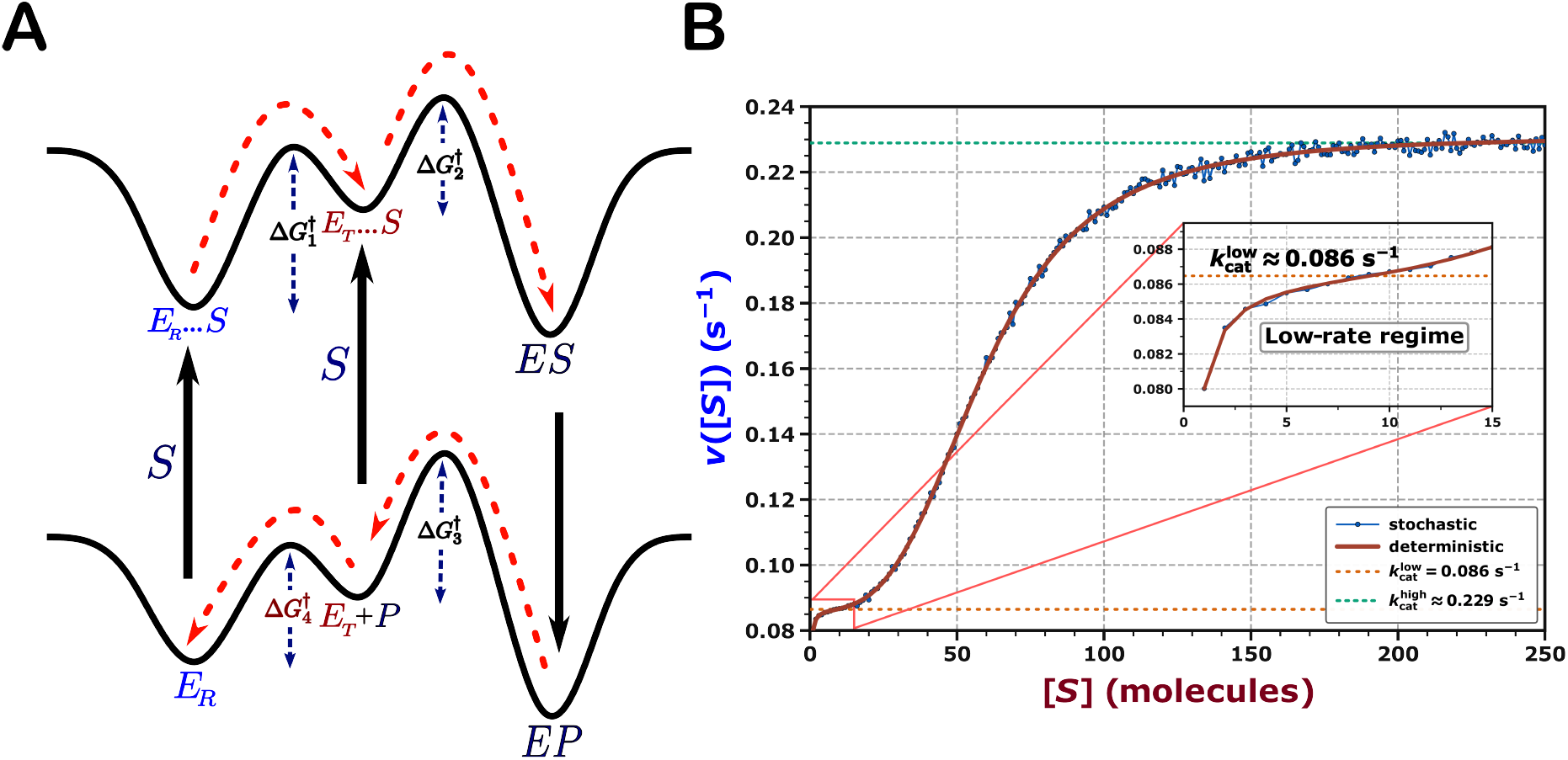
Free-energy landscape and substrate-dependent switching between turnover regimes. **(A)** Competing substrate-entry routes and associated activation barriers in the extended catalytic network. **(B)** Steady-state velocity, *v*([*S*]), showing close agreement between stochastic simulations and the deterministic model and a crossover between the low- and high-turnover plateaus; the inset highlights the low-rate saturation regime.

Mechanistically, the catalytic cycle can be initiated from either conformational state. Starting from the resting conformation (*E*_*R*_), substrate association initially yields a transient encounter complex (*E*_*R*_ … *S*), which must surmount the activation barrier 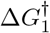 to undergo a structural transition into the intermediate state (*E*_*T*_ … *S*). Alternatively, the enzyme primed in its high-affinity conformation (*E*_*T*_) can directly capture the substrate to converge upon the identical *E*_*T*_ … *S* intermediate. From this shared on-pathway species, the system undergoes structural stabilization across the barrier 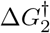 to form the stable enzyme–substrate (*ES*) complex. The system then transitions from the substrate surface to the product free-energy surface, driving the transformation of the *ES* complex into the enzyme–product (*EP*) complex. Subsequent product release, governed by the activation barrier 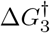, regenerates the free *E*_*T*_ conformation, thereby returning the enzyme to a critical kinetic bifurcation point. Here, the system must either overcome the relaxation barrier 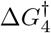 to return to the *E*_*R*_ state or immediately engage another substrate molecule to sustain high-velocity turnover. The free-energy landscape integrates parallel entry routes into a convergent pathway, where the activation barrier 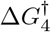 determines whether the enzyme resets to the slower catalytic route or remains primed for the faster pathway.

This kinetic scheme can be rigorously analyzed using complementary stochastic and deterministic modeling frameworks, both capturing equivalent aspects of the enzyme kinetics through distinct modeling approaches. Within the stochastic framework, the system is modeled as a discrete-state, continuous-time Markov process where transitions between conformational and chemical states occur probabilistically based on their microscopic rate constants. To evaluate the catalytic velocity as a function of substrate concentration, the Gillespie algorithm is employed iteratively across a discrete spectrum of fixed [*S*] values. For each fixed substrate copy number *S*, independent trajectories were simulated until a predefined number of product-release events occurred. The waiting times between consecutive turnovers were averaged to determine the mean first-passage time, *τ*_SSA_(*S*) = ⟨Δ*t*_*P*_⟩. The steady-state velocity was subsequently calculated using

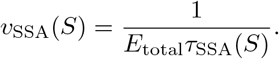

For a single-enzyme system (*E*_total_ = 1), this expression simplifies to

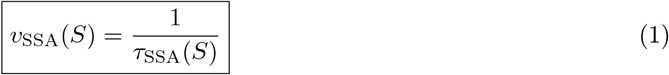

Complementing the stochastic simulations, the corresponding deterministic rate law is derived analytically by encoding the identical network of state transitions into a generator matrix *Q*. Solving the associated linear system yields the exact mean first-passage time (MFPT) from the initial enzyme state to the absorbing product state. This first-passage formalism provides a closed-form expression for the overall turnover time *τ*_overall_:

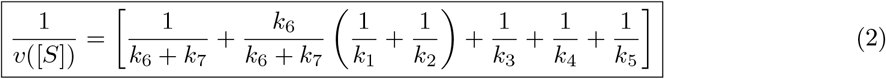

where the steady-state catalytic velocity *v*([*S*]) emerges naturally as the reciprocal of the total transition time. Comprehensive mathematical derivations, including the explicit *Q*-matrix construction and step-by-step inversion procedures, are detailed in the Supporting Information (S1).

## Results & Discussion

### Distinct Turnover Regimes Emerge from Barrier-Controlled Switching

The complete kinetic scheme shown in Fig. 2A is simulated using the Gillespie algorithm, with the stochastic catalytic velocity calculated according to Eq. 1. Alongside these simulations, the deterministic steady-state rate expression defined in Eq. 2 is evaluated across the same range. Fig. 2B displays the catalytic velocity *v*([*S*]) against substrate concentration [*S*] using the parameters from Table 1, where the blue dots represent discrete data points from the stochastic simulations and the solid red line connects them using the deterministic rate profile. The two approaches show excellent agreement across the entire substrate range. At low substrate amounts, the enzyme stays mostly in the slow pathway, resulting in a low baseline rate of 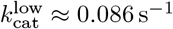. As [*S*] increases, more frequent substrate-binding events drive a smooth transition, eventually leveling off at a higher maximum rate of 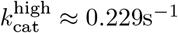.

**Table 1.** Elementary reactions, rate laws, and corresponding parameters.

| Elementary reaction | Rate law | Parameters |
| --- | --- | --- |
| $E_R + S \rightarrow E_R \cdots S$ | $k_1 = 4\pi D_s a_s [S]$ | $4\pi D_s a_s = 1.0$ |
| $E_R \cdots S \rightarrow E_T \cdots S$ | $k_2 = A_1 \exp[-\Delta G_1^\ddagger / (k_B T)]$ | $A_1 = 1, \Delta G_1^\ddagger = 1.5 k_B T$ |
| $E_T \cdots S \rightarrow ES$ | $k_3 = A_2 \exp[-\Delta G_2^\ddagger / (k_B T)]$ | $A_2 = 1, \Delta G_2^\ddagger = 0.5 k_B T$ |
| $ES \rightarrow EP$ | $k_4 = \text{constant}$ | $k_4 = 1.0$ |
| $EP \rightarrow E_T + P$ | $k_5 = A_3 \exp[-\Delta G_3^\ddagger / (k_B T)]$ | $A_3 = 1, \Delta G_3^\ddagger = 0.5 k_B T$ |
| $E_T \rightarrow E_R$ | $k_6 = A_4 \exp[-\Delta G_4^\ddagger / (k_B T)]$ | $A_4 = 1, \Delta G_4^\ddagger = 1.0 k_B T$ |
| $E_T + S \rightarrow E_T \cdots S$ | $k_7 = k_7^{\max} \frac{[S]^n}{K^n + [S]^n}$ | $k_7^{\max} = 75, K = 225, n = 3.25$ |

Mechanistically, product formation can proceed through two distinct catalytic routes. The balance between these pathways depends on a key kinetic bottleneck governed by the relative timescales of substrate capture (*E*_*T*_ → *E*_*T*_ … *S*) versus conformational relaxation (*E*_*T*_ → *E*_*R*_) over the critical barrier 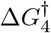. At low substrate copy numbers, the system tends to relax, forcing the enzyme to traverse the longer catalytic cycle and resulting in the lower rate threshold. In contrast, at higher substrate availability, rapid substrate capture makes the *E*_*T*_ → *E*_*T*_ … *S* transition much faster. The enzyme bypasses the relaxation step and follows the shorter catalytic route, allowing the system to reach the upper saturation plateau. Consequently, the enzyme stays in a low-velocity regime below a critical substrate concentration and transitions into a high-velocity regime above it. The kinetic switching between the two distinct catalytic regimes is evident in Fig. 2B, with the inset providing an expanded view of the low-rate regime, showing its saturation near 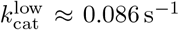. To systematically dissect the mechanistic origin of this bimodal kinetic behaviour, the complete landscape shown in Fig. 2A can be partitioned into two independent, topologically isolated sub-cycles:

- **The Slow Catalytic Route:** Eliminating the direct binding pathway from the tensed state forces the enzyme to exclusively traverse the extended catalytic loop, yielding the modified free energy landscape shown in Fig. S1(A). Under these conditions, every turnover must overcome the high conformational relaxation barrier 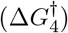 to reset the enzyme. Lacking the accelerated alternative route, the system exhibits no kinetic switching and flattens into a uniform low-velocity profile with a baseline rate of 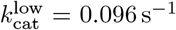, with both the deterministic and stochastic rate profiles projected together in Fig. S1(B).
- **The Faster Catalytic Cycle:** Conversely, omitting the relaxed state (*E*_*R*_) and its associated slow recovery steps forces substrate capture to occur entirely via the tensed conformation (*E*_*T*_), yielding the modified free energy landscape shown in Fig. S2(A). This architecture isolates a streamlined, high-efficiency catalytic loop that bypasses the relaxation bottleneck completely. Consequently, the enzyme operates continuously within its upper kinetic limit, causing the steady-state velocity to level off at a significantly elevated maximum rate of 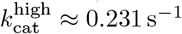, with both the deterministic and stochastic rate profiles projected together in Fig. S2(B).

The limiting turnover rates of the isolated slow and fast subcycles closely match the lower and upper plateaus in Fig. 2B, confirming that the two plateaus originate from the distinct kinetic limits of these competing catalytic routes.

### Barrier Dependence of the Kinetic Crossover

The role of the conformational relaxation barrier 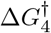, which mediates the transition between the tensed (*E*_*T*_) and relaxed (*E*_*R*_) states, was examined by systematically varying its value from 0.5 *k*_B_*T* to 5.0 *k*_B_*T* in increments of 0.5 *k*_B_*T* . This generates ten distinct velocity–substrate profiles in both the stochastic and deterministic frameworks. Altering the barrier height primarily modulates the catalytic rate at low and intermediate substrate levels, whereas the profiles converge toward the same high-substrate saturation plateau. Within the present kinetic model, the resulting curves also pass through a well-defined common crossover region, giving rise to an isosbestic-like kinetic response.

Figure 3A shows the ensemble of rate–substrate profiles obtained from stochastic simulations, while Fig. 3B presents the corresponding deterministic results. Both descriptions reproduce a closely defined common intersection. The stochastic simulations locate this crossover at approximately *S*_iso_ ≈ 37.7 molecules and *v*_iso_ ≈ 0.114 s^−1^, while the deterministic model gives a closely matching coordinate of *S*_iso_ ≈ 38.0 molecules and *v*_iso_ ≈ 0.115 s^−1^. The dependence of this crossover on 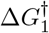 was examined further in Supplementary Figs. S3 and S4. For each fixed value of 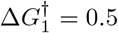, 1.0, 1.5, and 2.0 *k*_B_*T* , the barrier 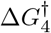 was varied over the same range. In both the stochastic and deterministic calculations, the corresponding rate–substrate profiles retain a well-defined common crossing. As 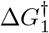 increases, the position of this crossing shifts toward lower substrate copy number and lower catalytic velocity. This behavior reflects the role of 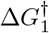 in regulating the *E*_*R*_ … *S E*_*T*_ … *S* transition that connects the two catalytic cycles. A larger barrier slows this conversion and increases the residence of the enzyme in the *E*_*R*_ … *S* intermediate associated with the slower catalytic loop, thereby shifting the crossover toward lower substrate levels and lower turnover rates. Conversely, reducing the barrier facilitates transfer into the *E*_*T*_ … *S* intermediate of the faster catalytic loop, shifting the crossover toward higher substrate levels and higher catalytic velocities.

**Figure 3.**
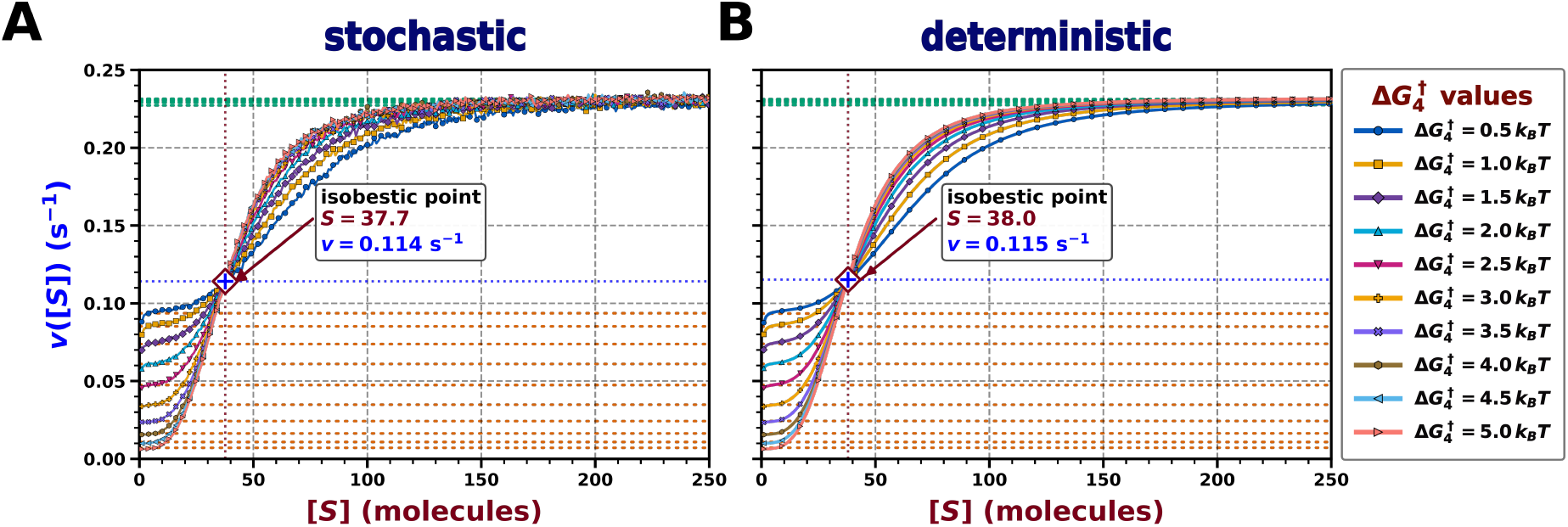
Barrier-dependent steady-state kinetics under systematic variation of the conformational relaxation barrier 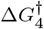 from 0.5 to 5.0 *k*_B_*T* . **(A)** Stochastic and **(B)** deterministic rate–substrate profiles. Both frameworks exhibit a common crossover region while approaching the same saturation plateau at high substrate levels.

Supplementary Fig. S5 summarizes this shift in terms of the characteristic crossover velocity, *v*_*c*_ = *v*(*S*_*c*_), and the corresponding substrate value, *S*_*c*_. Over the parameter range examined, *v*_*c*_ varies approximately linearly with 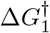, whereas *S*_*c*_ follows an approximately exponential decay.

### Kinetic Partitioning of Catalytic Flux between Fast and Slow Turnover Pathways

The extended free-energy landscape contains two distinct conformational transitions. The barrier 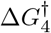 governs interconversion between the unbound conformations *E*_*T*_ and *E*_*R*_, whereas the parallel barrier 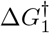 controls the corresponding transition between the weakly substrate-bound states, *E*_*R*_ … *S* and *E*_*T*_ … *S*. We therefore examine 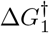 separately to determine how conformational exchange within the substrate-bound ensemble influences catalytic progression, thereby regulating catalytic flux and the attainable turnover rate.

The barrier 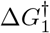 was varied from 0.5 *k*_B_*T* to 5.0 *k*_B_*T* over ten equally spaced values. Fig. 4A presents the stochastic rate–substrate profiles obtained for these barrier heights. A systematic decrease in the catalytic velocity is observed as 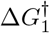 increases, with the separation between the curves becoming particularly pronounced in the high-substrate regime. Fig. 4B shows the corresponding rate profiles calculated from the deterministic expression in Eq. 2. The analytical results reproduce the same barrier-dependent reduction in catalytic velocity and the progressive lowering of the upper saturation plateau. The macroscopic kinetic suppression and saturation plateau depression observed under 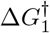 modulation are mechanistically rationalized by partitioning the microscopic turnover trajectories into discrete fast and slow catalytic cycles. The fast cycle follows *E*_*T*_ +*S* → *E*_*T*_ … *S* → *ES* → *EP* → *E*_*T*_ +*P* and therefore bypasses the 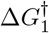 -dependent conformational transition. In contrast, the slow cycle follows *E*_*T*_ → *E*_*R*_, followed by *E*_*R*_ + *S* → *E*_*R*_ … *S* → *E*_*T*_ … *S* → *ES* → *EP* → *E*_*T*_ + *P* . This route contains the additional conformational relaxation and substrate-binding steps, as well as the barrier-controlled conversion of *E*_*R*_ … *S* into *E*_*T*_ … *S*. At *S* = 1000, the fractions of turnover events following the two routes are

**Figure 4.**
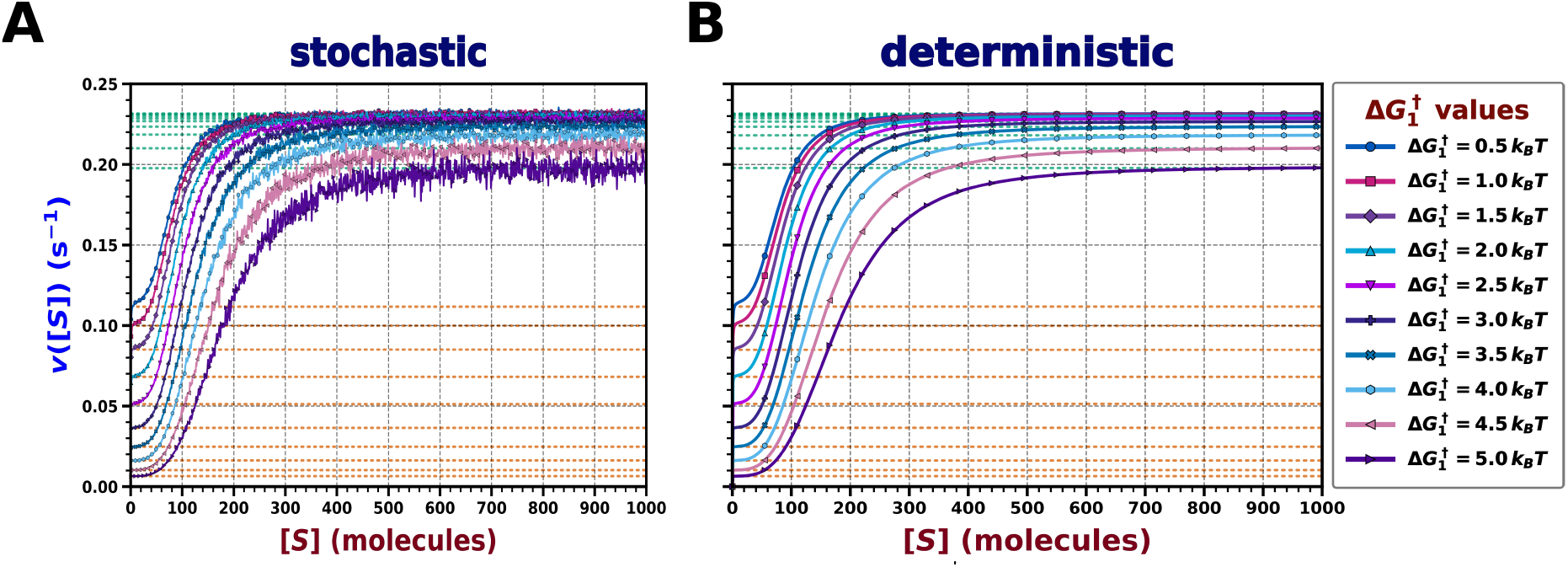
Influence of the substrate-bound conformational barrier 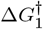 on steady-state kinetics. **(A)** Stochastic and **(B)** deterministic rate–substrate profiles mapped across a range of 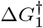 values from 0.5 to 5.0 *k*_B_*T* . Incremental elevation of the activation barrier induces a systematic attenuation of the reaction velocity and a pronounced depression of the asymptotic saturation plateau.

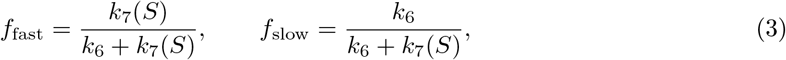

which yield *f*_fast_ = 0.9951 and *f*_slow_ = 0.0049. The overall mean first-passage time and the corresponding turnover rate are given by

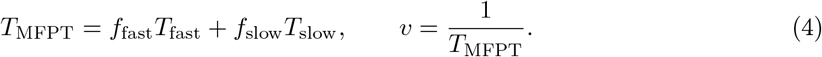

For the lowest barrier, 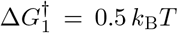, the fast-and slow-cycle MFPTs are 4.3108 s and 5.9605 s, respectively. Substitution of these values and the route fractions from Eq. 3 into Eq. 4 gives *T*_MFPT_ = 4.3189 s and *v* = 0.2315 s^−1^, with the slow cycle contributing approximately 0.68% of the total turnover time. At the highest barrier, 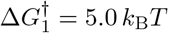 , the fast-cycle MFPT remains 4.3108 s, whereas the slow-cycle MFPT increases to 152.7250 s. Consequently, Eq. 4 gives *T*_MFPT_ = 5.0409 s and *v* = 0.1984 s^−1^, while the slow-cycle contribution to the total turnover time increases to approximately 14.9%. Thus, despite its small probability of occurrence, the greatly prolonged slow cycle at large 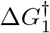 increases the overall MFPT and reduces the attainable catalytic rate.

### Barrier-Controlled Emergence of Burst–Halt Catalytic Turnover

The preceding MFPT analysis establishes that the reduction in catalytic turnover arises from occasional entry into the slow catalytic cycle, whose duration increases strongly with 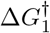. To examine how this mechanism appears in individual trajectories, we consider the cumulative product count *P* (*t*) at fixed substrate concentration. Unlike the time-averaged turnover rate, *P* (*t*) preserves the temporal sequence of successive product-release events, allowing periods of sustained catalytic activity to be distinguished from prolonged pauses associated with residence in the slow cycle.

Experimentally, single-enzyme turnover has been shown to proceed intermittently, with periods of rapid product formation separated by extended intervals of low catalytic activity, particularly at high substrate concentration. Xie and co-workers demonstrated this behavior using immobilized single *β*-galactosidase molecules and a fluorogenic substrate, such that each turnover was detected as an individual fluorescent product burst. At low substrate concentration, the waiting times between successive turnovers were approximately monoexponential, consistent with substrate binding as the dominant kinetic limitation. At higher substrate concentration, however, the turnover events became temporally clustered, producing broad, multiexponential waiting-time distributions and correlations between successive events. [53] A complementary trajectory-based Gillespie analysis was performed at selected high substrate levels to characterize burst–halt kinetics, consistent with the experimentally observed temporal clustering of turnover events under high-substrate conditions discussed earlier. This analysis was carried out separately from the steady-state calculations used to construct the rate–substrate profiles described above. In the earlier approach, the waiting times between successive product-release events were averaged at each fixed *S* to obtain a single turnover rate, *v*(*S*). Here, the complete product-release history was retained and the cumulative product count, *P* (*t*), was followed along one long trajectory. Simulations were performed at *S* = 250, 500, 750, and 1000, while 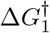 was varied from 1 to 6 *k*_B_*T* in steps of 1 *k*_B_*T* . Fig. 5A–D show the corresponding trajectories. At smaller barrier heights, *P* (*t*) increases nearly steadily, indicating regular product release. At 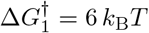 , rapid product formation is interrupted by clear plateaus, with the burst–halt pattern most evident at *S* = 750 and *S* = 1000. These pauses arise from prolonged residence in the slow catalytic cycle and reduce the average slope of *P* (*t*), thereby lowering the effective turnover rate.

**Figure 5.**
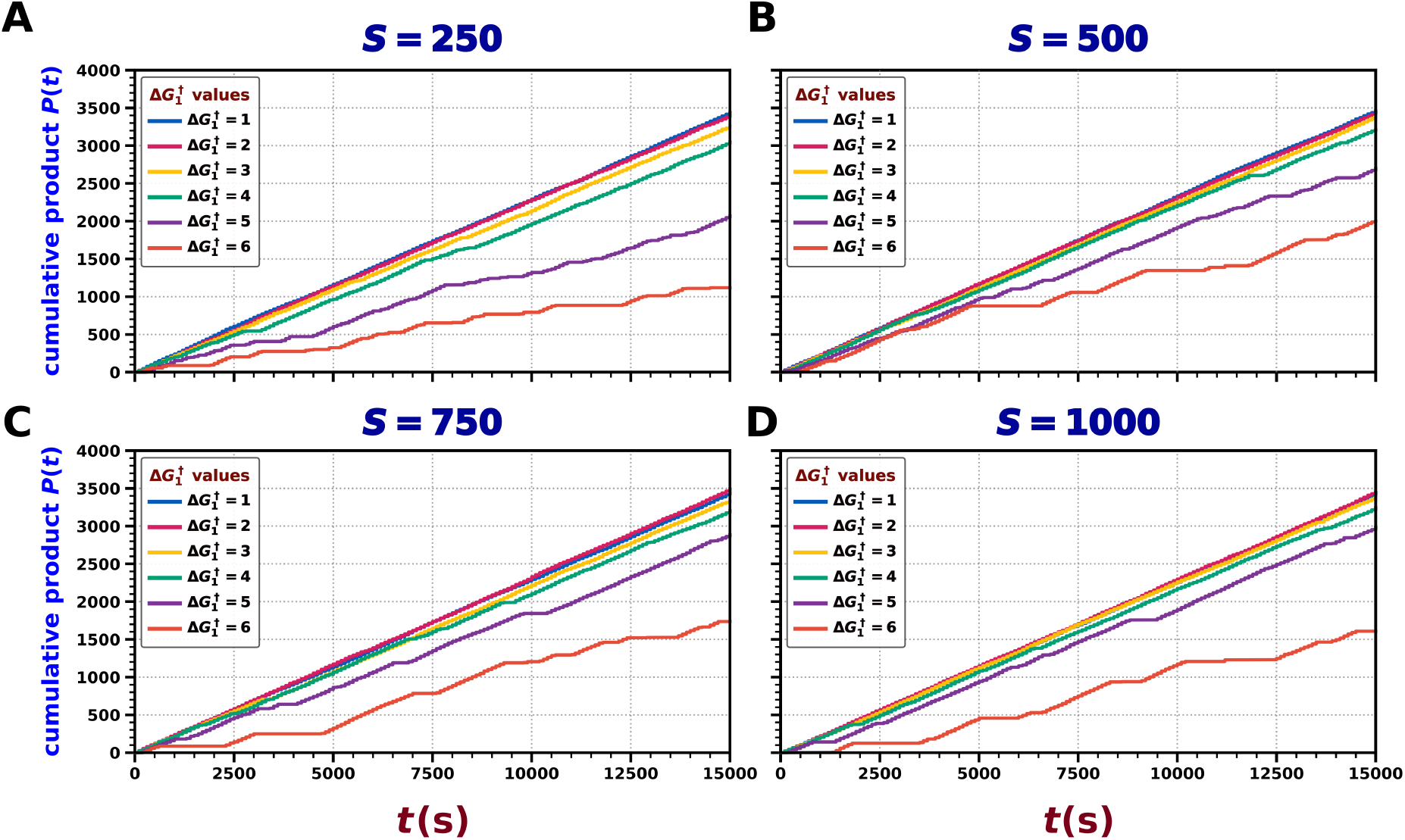
Single-trajectory resolution of product-release dynamics showing barrier-induced temporal intermittency. Stochastic trajectories of cumulative product generation, *P* (*t*), were modeled via Gillespie simulations at fixed substrate concentrations: **(A)** *S* = 250, **(B)** *S* = 500, **(C)** *S* = 750, and **(D)** *S* = 1000. Under low conformational activation barriers 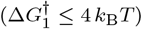, catalytic turnover is quasi-continuous, yielding linear production profiles. In contrast, elevating the barrier to 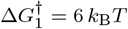 induces pronounced burst–halt kinetics, marked by extended macroscopic pauses (plateaus) that become increasingly distinct at high substrate concentrations (*S ≥* 750).

Supplementary Fig. S6 compares the effects of the two conformational barriers at *S* = 500. In panel A, 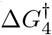 is fixed at 1.0 *k*_B_*T* , while 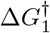 is varied from 0.5 to 5.0 *k*_B_*T* ; the largest barrier produces clear plateaus in *P* (*t*), consistent with the burst–halt behavior discussed above. In panel B, 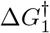 is fixed at 1.5 *k*_B_*T* , while 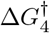 is varied over the same range. These trajectories remain nearly linear, indicating that varying 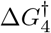 does not produce the same separation between the fast- and slow-cycle timescales or the corresponding pauses in product release. Thus, varying 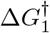 separates the fast and slow catalytic timescales sufficiently to produce distinct intermittent turnover, whereas varying 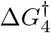 mainly changes the pathway selection without creating the same long-lived interruption in product formation. Together, these results show a clear correspondence between the intermittent turnover observed experimentally and the burst–halt behavior obtained from the present model. In both cases, periods of rapid product formation are separated by long inactive intervals arising from slow conformational transitions.

### Canonical Kinetic Signatures Emerging from a Bimodal Catalytic Landscape

The extended kinetic scheme can recover qualitatively distinct canonical rate–substrate responses depending on the relative accessibility of the two catalytic pathways. At the limiting values of the conformational relaxation barrier 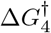, the stochastic profiles become dominated by one of the two routes and, over the substrate range examined, approach an effectively single-saturation response reminiscent of classical Michaelis–Menten kinetics.

In Fig. 6A, the large relaxation barrier 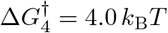 suppresses the *E*_*T*_ → *E*_*R*_ transition and retains most of the catalytic flux within the faster *E*_*T*_ -initiated pathway. Using 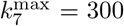, *K* = 3500, and *n* = 1.2, with *k*_2_ = *e*^−2.5^, the rate increases monotonically and approaches the upper turnover limit, 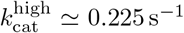. Conversely, Fig. 6B represents the small-barrier limit, 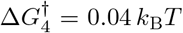 , for which rapid relaxation toward *E*_*R*_ strongly favors the slower catalytic route. For 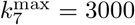, *K* = 500, and *n* = 20, while retaining *k*_2_ = *e*^−2.5^, the stochastic profile again appears as a single saturating response, but now approaches the lower turnover limit, 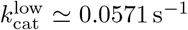. Thus, preferential selection of either catalytic pathway produces Michaelis–Menten-like hyperbolic saturation at the corresponding lower or upper kinetic limit.

**Figure 6.**
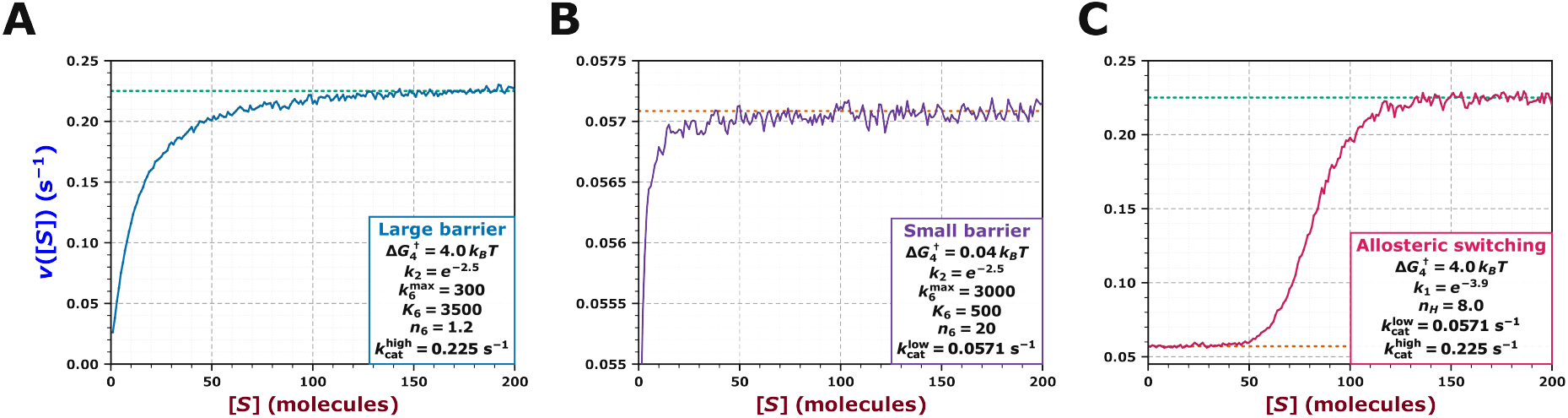
Michaelis–Menten-like saturation at limiting conformational barriers. **(A)** At 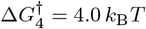 , the stochastic rate approaches the higher saturation limit, 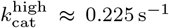. **(B)** At 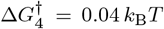 , the rate approaches the lower saturation limit, 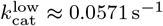. The allosteric-like kinetic signature is illustrated in **(C)**, where the stochastic response undergoes a sigmoidal transition from the lower saturation limit, 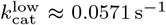, to the higher saturation limit, 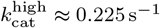.

A different limiting response is obtained when the substrate dependence of pathway selection is made sufficiently sharp. As shown in Fig. 6C, the turnover rate then undergoes a pronounced sigmoidal transition between the lower and upper kinetic regimes. The resulting profile closely resembles the cooperative response conventionally associated with allosteric enzyme kinetics, with the low- and high-rate limits remaining approximately 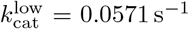 and 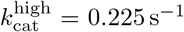, respectively. The same underlying kinetic architecture can therefore generate either an apparently hyperbolic or sigmoidal substrate response through modulation of conformational pathway selection.

Importantly, the apparent Michaelis–Menten-like and allosteric-like responses do not eliminate the underlying bimodal character of the kinetic scheme. This becomes evident when the deterministic profiles are extended beyond the substrate window considered in the stochastic analysis. As shown in Supplementary Fig. S7(A), the large-barrier condition reveals an additional low-rate saturation regime at very low substrate levels, whereas Supplementary Fig. S7(B) shows that, under the small-barrier condition, the system ultimately crosses from the apparent low-rate plateau to the higher turnover regime at sufficiently large substrate levels. Thus, kinetic profiles that appear effectively single-saturating over a restricted substrate range may still retain signatures of the underlying low- and high-rate regimes. An analogous limitation may arise in experimental kinetic analyses, where rate–substrstrate profiles are typically interpreted within established kinetic formalisms over a finite range of accessible concentrations. If an additional turnover regime emerges only at the extreme low- or high-substrate limits, it may remain experimentally unresolved and thereby escape detection within the concentration range used to characterize the apparent kinetic response.

## Discussion and Conclusions

The present study extends the classical Michaelis–Menten description from a free-energy landscape perspective by introducing a kinetic framework in which turnover can proceed through alternative conformational routes. Instead of assuming one dominant catalytic pathway with a single saturation limit, the enzyme redistributes flux between a slower and a faster route depending on substrate availability and conformational barrier heights. In this landscape, direct substrate binding to the pre-existing tensed state represents a conformational-selection-like route, because the substrate is captured by a state that is already closer to the productive catalytic configuration. In contrast, substrate binding to the relaxed state followed by a substrate-bound conformational transition represents an induced-fit-like route, because productive catalysis requires an additional rearrangement after binding. The stochastic simulations and deterministic mean-first-passage-time analysis show that competition between these two routes gives rise to distinct lower and higher turnover regimes connected by a substrate-dependent transition. At lower substrate levels, relaxation toward the relaxed state favours the longer induced-fit-like route and produces a lower effective rate. As substrate availability increases, direct capture by the tensed state becomes more probable, shifting the flux toward the shorter conformational-selection-like route and producing a higher turnover regime. The observed kinetic response therefore reflects a changing balance between conformational selection, induced rearrangement, and catalytic progression, rather than a single Michaelis–Menten pathway.

The two conformational barriers have distinct kinetic roles. The relaxation barrier mainly controls how catalytic flux is divided between the two routes, and its modulation gives rise to the common crossing of the rate–substrate profiles. The agreement between the stochastic and deterministic descriptions shows that this kinetic isosbestic point arises from the structure of the reaction scheme rather than from simulation noise. In contrast, the substrate-bound conformational barrier primarily controls the duration of the slow route. Increasing this barrier prolongs residence in the weakly bound intermediate, increases the total turnover time, and lowers the attainable saturation rate. The MFPT analysis further shows that even a rarely visited slow pathway can strongly influence the measured rate when its completion time is much longer than that of the fast pathway.

The same separation of timescales is visible when cumulative product release is followed along a single long trajectory at a fixed substrate level. At smaller barriers, successive product-release events are distributed almost uniformly in time, whereas larger barriers produce clusters of product formation separated by extended pauses. This burst–halt behaviour closely resembles single-molecule observations for *β*-galactosidase, cholesterol oxidase, and lipase, where periods of high catalytic activity are separated by low-activity intervals and accompanied by broad or correlated turnover times. These experimental features have been associated with slow conformational fluctuations that change catalytic activity over time. The agreement with the present fixed-*S* trajectories therefore supports the interpretation that switching between kinetically distinct routes can affect both individual turnover patterns and the time-averaged catalytic rate.

At limiting conditions, the same kinetic framework can approach either an effectively single-saturating Michaelis–Menten-like response or a sigmoidal allosteric-like response, depending on how conformational pathway selection is modulated. Importantly, these apparent kinetic limits do not eliminate the underlying bimodal architecture; when explored over a sufficiently broad substrate range, the distinct low- and high-turnover regimes remain recoverable within the same conformational landscape. From an experimental perspective, enzyme rate–substrate profiles are commonly characterized over a finite concentration window, within which a hyperbolic dependence is typically interpreted as Michaelis– Menten-like kinetics, whereas a sigmoidal profile is taken to indicate an allosteric or cooperative response. However, such phenomenological descriptions are necessarily conditioned by the substrate range that is experimentally accessible. If measurements sample only a limited portion of the kinetic landscape, alternative catalytic routes may remain unresolved. Extending the substrate range toward very low or high concentrations can reveal these hidden regimes, although such conditions may be difficult to probe experimentally. The present framework therefore suggests that apparently distinct kinetic forms may correspond to different observable regimes of the same underlying conformationally coupled reaction landscape.

Taken together, these results identify conformationally controlled flux redistribution as a key factor shaping the observed kinetic behavior. Changes in barrier crossing, pathway occupancy, and the time spent within each catalytic route can influence both steady-state turnover and the temporal organization of individual catalytic events. These changes can give rise to distinct saturation regimes, kinetic crossovers, and burst–halt dynamics within the same reaction landscape. The present formulation should therefore be regarded as a reduced mechanistic representation that captures the essential kinetic consequences of conformational exchange without attempting to reproduce the full complexity of enzymatic dynamics. In this context, the present framework provides a physically interpretable basis for relating conformational rearrangements and pathway redistribution to the diverse kinetic responses observed across enzyme-catalyzed reactions. The framework thus offers a mechanistic description of more complex catalytic processes in which multiple conformational states and competing pathways shape experimentally observed kinetics and biologically relevant enzyme function.

## Acknowledgments

The authors sincerely thank the Central Supercomputing Facility of the Indian Association for the Cultivation of Science, Kolkata and ANRF (ANRF grant ANRF/ARG/2025/003888/CS) for providing essential computational resources. Rohon Mitra gratefully acknowledges the Council for Scientific and Industrial Research (CSIR) for the award of a research fellowship, which supported this work.

## References

[1] Geoffrey M. Cooper. The central role of enzymes as biological catalysts. In The Cell: A Molecular Approach. Sinauer Associates, Sunderland, MA, 2 edition, 2000.

[2] Peter K. Robinson. Enzymes: principles and biotechnological applications. Essays in Biochemistry, 59:1, 2015.

[3] Thomas C. Bruice and Stephen J. Benkovic. Chemical basis for enzyme catalysis. Biochemistry, 39(21):6267–6274, 2000.

[4] John P. Richard. Enzymatic rate enhancements: A review and perspective. Biochemistry, 52(12):2009–2011, 2013.

[5] Stephen J. Benkovic and Sharon Hammes-Schiffer. A perspective on enzyme catalysis. Science, 301(5637):1196–1202, 2003.

[6] Pratul K. Agarwal. A biophysical perspective on enzyme catalysis. Biochemistry, 58(6):438, 2018.

[7] Robert Callender and R. Brian Dyer. The dynamical nature of enzymatic catalysis. Accounts of Chemical Research, 48(2):407–413, 2015.

[8] Arieh Warshel and Ram Prasad Bora. Perspective: Defining and quantifying the role of dynamics in enzyme catalysis. Journal of Chemical Physics, 144(18), 2016.

[9] Kenneth A. Johnson. Fitting enzyme kinetic data with KinTek Global Kinetic Explorer. In Methods in Enzymology, volume 467, pages 601–626. Elsevier, 2009.

[10] Elan Zohar Eisenmesser, Daryl A. Bosco, Mikael Akke, and Dorothee Kern. Enzyme dynamics during catalysis. Science, 295(5559):1520–1523, 2002.

[11] Elan Z. Eisenmesser, Oscar Millet, Wladimir Labeikovsky, Dmitry M. Korzhnev, Magnus Wolf-Watz, Daryl A. Bosco, Jack J. Skalicky, Lewis E. Kay, and Dorothee Kern. Intrinsic dynamics of an enzyme underlies catalysis. Nature, 438(7064):117–121, 2005.

[12] Katherine Henzler-Wildman and Dorothee Kern. Dynamic personalities of proteins. Nature 2007 450:7172, 450(7172):964–972, 2007.

[13] H. A. Kramers. Brownian motion in a field of force and the diffusion model of chemical reactions. Physica, 7(4):284–304, 1940.

[14] Rohon Mitra and Biman Jana. A model of protein folding with multiple native states: Metamorphicity, intrinsic disorderness, and folding upon binding of proteins. Journal of Chemical Physics, 163, 8 2025.

[15] Bruce J. Berne, Michal Borkovec, and John E. Straub. Classical and modern methods in reaction rate theory. Journal of Physical Chemistry, 92(13):3711–3725, 1988.

[16] Hans Frauenfelder, Stephen G. Sligar, and Peter G. Wolynes. The energy landscapes and motions of proteins. Science, 254(5038):1598–1603, 1991.

[17] George Edward Briggs and John Burdon Sanderson Haldane. A note on the kinetics of enzyme action. The Biochemical journal, 19(2):338–339, 1925.

[18] Kenneth A. Johnson and Roger S. Goody. The original michaelis constant: Translation of the 1913 michaelis-menten paper. Biochemistry, 50(39):8264, 2011.

[19] Athel Cornish-Bowden. The origins of enzyme kinetics. FEBS Letters, 587(17):2725–2730, 2013.

[20] Peter Hänggi, Peter Talkner, and Michal Borkovec. Reaction-rate theory: Fifty years after Kramers. Reviews of Modern Physics, 62(2):251–341, 1990.

[21] Rohon Mitra and Biman Jana. Local cooperative interactions reshape the folding transition in a one-dimensional spin-glass model. The Journal of Physical Chemistry B, 8 2026.

[22] D. E. Koshland. Application of a theory of enzyme specificity to protein synthesis. Proceedings of the National Academy of Sciences, 44(2):98–104, 1958.

[23] Buyong Ma, Sandeep Kumar, Chung Jung Tsai, and Ruth Nussinov. Folding funnels and binding mechanisms. Protein Engineering, 12(9):713–720, 1999.

[24] Thomas R. Weikl and Carola Von Deuster. Selected-fit versus induced-fit protein binding: Kinetic differences and mutational analysis. Proteins: Structure, Function and Bioinformatics, 75(1):104–110, 2009.

[25] Austin D. Vogt and Enrico Di Cera. Conformational selection is a dominant mechanism of ligand binding. Biochemistry, 52(34):5723–5729, 2013.

[26] Gordon G. Hammes, Yu Chu Chang, and Terrence G. Oas. Conformational selection or induced fit: A flux description of reaction mechanism. Proceedings of the National Academy of Sciences of the United States of America, 106(33):13737–13741, 2009.

[27] Stefano Gianni, Jakob Dogan, and Per Jemth. Distinguishing induced fit from conformational selection. Biophysical Chemistry, 189:33–39, 2014.

[28] Fabian Paul and Thomas R. Weikl. How to distinguish conformational selection and induced fit based on chemical relaxation rates. PLOS Computational Biology, 12(9):e1005067, 2016.

[29] Sarah M. Sullivan and Todd Holyoak. Enzymes with lid-gated active sites must operate by an induced fit mechanism instead of conformational selection. Proceedings of the National Academy of Sciences of the United States of America, 105(37):13829–13834, 2008.

[30] Jacque Monod, Jeffries Wyman, and Jean Pierre Changeux. On the nature of allosteric transitions: A plausible model. Journal of Molecular Biology, 12(1):88–118, 1965.

[31] D. E. Koshland, J. G. Nemethy, and D. Filmer. Comparison of experimental binding data and theoretical models in proteins containing subunits. Biochemistry, 5(1):365–385, 1966.

[32] Jean Pierre Changeux and Stuart J. Edelstein. Allosteric mechanisms of signal transduction. Science, 308(5727):1424–1428, 2005.

[33] Jean Pierre Changeux. Allostery and the monod–wyman–changeux model after 50 years. Ann. Rev. Biophys., 41(1):103–133, 2012.

[34] C. Frieden. Slow transitions and hysteretic behavior in enzymes. Annual review of biochemistry, 48:471–489, 1979.

[35] J. Ricard. Generalized microscopic reversibility, kinetic co-operativity of enzymes and evolution. The Biochemical journal, 175(3):779–791, 1978.

[36] Jean-Claude-C MEUNIER, Jean BUC, André NAVARRO, and Jacques RECARD. Regulatory behavior of monomeric enzymes: 2. a wheat-germ hexokinase as a mnemonical enzyme. European Journal of Biochemistry, 49(1):209–223, 1974.

[37] Carol M. Porter and Brian G. Miller. Cooperativity in monomeric enzymes with single ligandbinding sites. Bioorganic Chemistry, 43:44–50, 2012.

[38] A. C. Storer and A. Cornish Bowden. Kinetics of rat liver glucokinase. cooperative interactions with glucose at physiologically significant concentrations. Biochemical Journal, 159(1):7–14, 1976.

[39] Maria Luz CÁRDENAS, Eliana RABAJILLE, and Hermann NIEMEYER. Suppression of kinetic cooperativity of hexokinase d (glucokinase) by competitive inhibitors: A slow transition model. European Journal of Biochemistry, 145(1):163–171, 1984.

[40] G. Pettersson. Mechanistic origin of the sigmoidal rate behaviour of glucokinase. Biochemical Journal, 233(2):347–350, 1986.

[41] Ofer Yifrach and Amnon Horovitz. Nested cooperativity in the atpase activity of the oligomeric chaperonin groel. Biochemistry, 34(16):5303–5308, 1995.

[42] Amnon Horovitz, Yael Fridmann, Galit Kafri, and Ofer Yifrach. Review: Allostery in chaperonins. Journal of Structural Biology, 135(2):104–114, 2001.

[43] Ranit Gruber and Amnon Horovitz. Allosteric mechanisms in chaperonin machines. Chemical Reviews, 116(11):6588–6606, 2016.

[44] Galit Kafri, Keith R. Willison, and Amnon Horovitz. Nested allosteric interactions in the cytoplasmic chaperonin containing tcp-1. Protein Science, 10(2):445–449, 2001.

[45] Galit Kafri and Amnon Horovitz. Transient kinetic analysis of atp-induced allosteric transitions in the eukaryotic chaperonin containing tcp-1. Journal of Molecular Biology, 326(4):981–987, 2003.

[46] David D. Boehr, Dan McElheny, H. Jane Dyson, and Peter E. Wrightt. The dynamic energy landscape of dihydrofolate reductase catalysis. Science, 313(5793):1638–1642, 2006.

[47] Karunesh Arora and Charles L. Brooks. Functionally important conformations of the met20 loop in dihydrofolate reductase are populated by rapid thermal fluctuations. Journal of the American Chemical Society, 131(15):5642, 2009.

[48] Gira Bhabha, Jeeyeon Lee, Damian C. Ekiert, Jongsik Gam, Ian A. Wilson, H. Jane Dyson, Stephen J. Benkovic, and Peter E. Wright. A dynamic knockout reveals that conformational fluctuations influence the chemical step of enzyme catalysis. Science, 332(6026):234–238, 2011.

[49] Saikat Dhibar and Biman Jana. Transferable collective variable to accelerate protein-ligand (un)binding transitions via explainable machine learning and intriguing role of ligand solvation. bioRxiv, page 2026.08.21.746233, 8 2026.

[50] H. P. Lu. Single-molecule enzymatic dynamics. Science, 282(5395):1877–1882, 1998.

[51] Noam Agmon. Conformational cycle of a single working enzyme. Journal of Physical Chemistry B, 104(32):7830–7834, 2000.

[52] X. Sunney Xie. Single-molecule approach to dispersed kinetics and dynamic disorder: Probing conformational fluctuation and enzymatic dynamics. The Journal of Chemical Physics, 117(24):11024–11032, 2002.

[53] Brian P. English, Wei Min, Antoine M. Van Oijen, Taek Lee Kang, Guobin Luo, Hongye Sun, Binny J. Cherayil, S. C. Kou, and X. Sunney Xie. Ever-fluctuating single enzyme molecules: Michaelis-menten equation revisited. Nature Chemical Biology, 2(2):87–94, 2006.

[54] Wei Min, Brian P. English, Guobin Luo, Binny J. Cherayil, S. C. Kou, and X. Sunney Xie. Fluctuating enzymes: lessons from single-molecule studies. Accounts of Chemical Research, 38(12):923–931, 2005.

[55] S. C. Kou, Binny J. Cherayil, Wei Min, Brian P. English, and X. Sunney Xie. Single-molecule michaelis - menten equations. Journal of Physical Chemistry B, 109(41):19068–19081, 2005.

[56] Hong Qian and Elliot L. Elson. Single-molecule enzymology: Stochastic michaelis-menten kinetics. Biophysical Chemistry, 101-102:565–576, 2002.

[57] Wei Min, X. Sunney Xie, and Biman Bagchi. Role of conformational dynamics in kinetics of an enzymatic cycle in a nonequilibrium steady state. The Journal of Chemical Physics, 131(6):065104, 2009.

[58] Vladi V. Heredia, Jim Thomson, David Nettleton, and Shaoxian Sun. Glucose-induced conformational changes in glucokinase mediate allosteric regulation: Transient kinetic analysis. Biochemistry, 45(24):7553–7562, 2006.

[59] Lars Edman, Zeno Földes-Papp, Stefan Wennmalm, and Rudolf Rigler. The fluctuating enzyme: A single molecule approach. Chemical Physics, 247(1):11–22, 1999.

[60] Vladimir Chernyak, Michael Schulz, and Shaul Mukamel. Stochastic-trajectories and nonpoisson kinetics in single-molecule spectroscopy. Journal of Chemical Physics, 111(16):7416–7425, 1999.

[61] George H. Weiss and Jaume Masoliver. Statistics of dwell times in a reaction with randomly fluctuating rates. Physica A: Statistical Mechanics and its Applications, 296(1–2):75–82, 2001.

[62] Jianshu Cao. Event-averaged measurements of single-molecule kinetics. Chemical Physics Letters, 327(1–2):38–44, 2000.

[63] Divya Singh, Tal Robin, Michael Urbakh, and Shlomi Reuveni. High-order michaelis-menten equations allow inference of hidden kinetic parameters in enzyme catalysis. Nature Communications, 16(1), 2025.

[64] Yann Sakref, Maitane Muñoz-Basagoiti, Zorana Zeravcic, and Olivier Rivoire. On kinetic constraints that catalysis imposes on elementary processes. Journal of Physical Chemistry B, 127(51):10950–10959, 2023.

[65] Asli Sahin, Daniel R. Weilandt, and Vassily Hatzimanikatis. Optimal enzyme utilization suggests that concentrations and thermodynamics determine binding mechanisms and enzyme saturations. Nature Communications, 14(1), 2023.

[66] S. Jordan Kerns, Roman V. Agafonov, Young Jin Cho, Francesco Pontiggia, Renee Otten, Dimitar V. Pachov, Steffen Kutter, Lien A. Phung, Padraig N. Murphy, Vu Thai, Tom Alber, Michael F. Hagan, and Dorothee Kern. The energy landscape of adenylate kinase during catalysis. Nature Structural and Molecular Biology, 22(2):124–131, 2015.

[67] Jingjing Guo and Huan Xiang Zhou. Protein allostery and conformational dynamics. Chemical Reviews, 116(11):6503–6515, 2016.

[68] Vincent J. Hilser, James O. Wrabl, and Hesam N. Motlagh. Structural and energetic basis of allostery. Ann. Rev. Biophys., 41(1):585–609, 2012.

[69] Hesam N. Motlagh, James O. Wrabl, Jing Li, and Vincent J. Hilser. The ensemble nature of allostery. Nature 2014 508:7496, 508(7496):331–339, 2014.

[70] Ruth Nussinov, Chung Jung Tsai, and Buyong Ma. The (still) underappreciated role of allostery in the cellular network. Annu. Rev. Biophys., 42(1):169–189, 2013.

[71] Tal Einav, Linas Mazutis, and Rob Phillips. Statistical mechanics of allosteric enzymes. Journal of Physical Chemistry B, 120(26):6021–6037, 2016.

