## Supporting Information for "Conformational Barrier-Driven Flux Redistribution across Competing Catalytic Pathways Unifies Diverse Enzyme Kinetic Regimes"

### 1 Supplementary Equations

#### S1: Derivation of the Deterministic Rate Expression from Elementary Steps

##### Population Balance Equations for the Kinetic Scheme

Since the elementary catalytic steps are already listed in the main manuscript, we begin directly from the population dynamics of the enzyme states. The state populations obey the following coupled kinetic equations:

$$\begin{aligned} \frac{d[E_R]}{dt} &= -k_1[E_R] + k_6[E_T], \\ \frac{d[E_R \cdots S]}{dt} &= k_1[E_R] - k_2[E_R \cdots S], \\ \frac{d[E_T]}{dt} &= k_5[EP] - (k_6 + k_7)[E_T], \\ \frac{d[E_T \cdots S]}{dt} &= k_2[E_R \cdots S] + k_7[E_T] - k_3[E_T \cdots S], \\ \frac{d[ES]}{dt} &= k_3[E_T \cdots S] - k_4[ES], \\ \frac{d[EP]}{dt} &= k_4[ES] - k_5[EP], \\ \frac{d[P]}{dt} &= k_5[EP]. \end{aligned} \tag{1}$$

The catalytic rate can be obtained by computing the mean first-passage time (MFPT) for one complete product-forming event. For this calculation, $E_T + P$ is treated as the absorbing product-released state.

##### Rate Matrix for the Transient States

We order the transient states as

$$\mathcal{S} = (E_T, E_R, E_R \cdots S, E_T \cdots S, ES, EP).$$

Using this ordering, the transient rate matrix is

$$\mathbf{Q} = \begin{pmatrix} -(k_6 + k_7) & k_6 & 0 & k_7 & 0 & 0 \\ 0 & -k_1 & k_1 & 0 & 0 & 0 \\ 0 & 0 & -k_2 & k_2 & 0 & 0 \\ 0 & 0 & 0 & -k_3 & k_3 & 0 \\ 0 & 0 & 0 & 0 & -k_4 & k_4 \\ 0 & 0 & 0 & 0 & 0 & -k_5 \end{pmatrix}. \tag{2}$$

The diagonal element of each row is the negative of the total rate of leaving that state, while the off-diagonal elements are the transition rates into the next accessible states.

##### MFPT Equations from the Rate Matrix

Let

$$\boldsymbol{\tau} = (\tau_{E_T}, \tau_{E_R}, \tau_{E_R \cdots S}, \tau_{E_T \cdots S}, \tau_{ES}, \tau_{EP})^T$$

be the vector of mean first-passage times to the absorbing product-released state. The backward MFPT equation is

$$-\mathbf{Q}\boldsymbol{\tau} = \mathbf{1}, \quad \mathbf{1} = (1, 1, 1, 1, 1, 1)^T. \tag{3}$$

Expanding Eq. (3) gives

$$(k_6 + k_7)\tau_{E_T} - k_6\tau_{E_R} - k_7\tau_{E_T \cdots S} = 1,$$

$$\begin{aligned} k_1\tau_{E_R} - k_1\tau_{E_R\cdots S} &= 1, \\ k_2\tau_{E_R\cdots S} - k_2\tau_{E_T\cdots S} &= 1, \\ k_3\tau_{E_T\cdots S} - k_3\tau_{ES} &= 1, \\ k_4\tau_{ES} - k_4\tau_{EP} &= 1, \quad k_5\tau_{EP} = 1. \end{aligned}$$

Solving backward from the product-releasing step gives

$$\begin{aligned} \tau_{EP} &= \frac{1}{k_5}, \\ \tau_{ES} &= \frac{1}{k_4} + \tau_{EP} = \frac{1}{k_4} + \frac{1}{k_5}, \\ \tau_{E_T\cdots S} &= \frac{1}{k_3} + \tau_{ES} = \frac{1}{k_3} + \frac{1}{k_4} + \frac{1}{k_5}, \\ \tau_{E_R\cdots S} &= \frac{1}{k_2} + \tau_{E_T\cdots S} = \frac{1}{k_2} + \frac{1}{k_3} + \frac{1}{k_4} + \frac{1}{k_5}, \\ \tau_{E_R} &= \frac{1}{k_1} + \tau_{E_R\cdots S} = \frac{1}{k_1} + \frac{1}{k_2} + \frac{1}{k_3} + \frac{1}{k_4} + \frac{1}{k_5}. \end{aligned} \tag{4}$$

##### Branching from $E_T$

From $E_T$, two exits are possible:

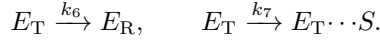

The total escape rate from $E_T$ is $k_6 + k_7$. Hence the mean waiting time in $E_T$ is $1/(k_6 + k_7)$, while the branching probabilities are

$$p_6 = \frac{k_6}{k_6 + k_7}, \quad p_7 = \frac{k_7}{k_6 + k_7}.$$

Therefore, the MFPT from $E_T$ is

$$\tau_{E_T} = \frac{1}{k_6 + k_7} + \frac{k_6}{k_6 + k_7}\tau_{E_R} + \frac{k_7}{k_6 + k_7}\tau_{E_T\cdots S}. \tag{5}$$

Substituting Eq. (4) into Eq. (5),

$$\tau_{E_T} = \frac{1}{k_6 + k_7} + \frac{k_6}{k_6 + k_7} \left( \frac{1}{k_1} + \frac{1}{k_2} + \frac{1}{k_3} + \frac{1}{k_4} + \frac{1}{k_5} \right) + \frac{k_7}{k_6 + k_7} \left( \frac{1}{k_3} + \frac{1}{k_4} + \frac{1}{k_5} \right).$$

The last three waiting-time terms are common to both branches. Therefore, using

$$\frac{k_6}{k_6 + k_7} + \frac{k_7}{k_6 + k_7} = 1,$$

we obtain the total turnover MFPT:

$$\tau_{\text{turn}}([S]) = \frac{1}{k_6 + k_7} + \frac{k_6}{k_6 + k_7} \left( \frac{1}{k_1} + \frac{1}{k_2} \right) + \frac{1}{k_3} + \frac{1}{k_4} + \frac{1}{k_5}. \tag{6}$$

##### Final Rate Expression

Since one product molecule is released after one completed cycle,

$$v([S]) = \frac{1}{\tau_{\text{turn}}([S])}.$$

Thus,

$$\boxed{\frac{1}{v([S])} = \frac{1}{k_6 + k_7} + \frac{k_6}{k_6 + k_7} \left( \frac{1}{k_1} + \frac{1}{k_2} \right) + \frac{1}{k_3} + \frac{1}{k_4} + \frac{1}{k_5}}. \tag{7}$$

With

$$k_1([S]) = 4\pi D_s a_s [S], \quad k_7([S]) = k_7^{\max} \frac{[S]^n}{K^n + [S]^n},$$

the explicit substrate-dependent form becomes

$$\frac{1}{v([S])} = \frac{1}{k_6 + k_7^{\max} \frac{[S]^n}{K^n + [S]^n}} + \frac{k_6}{k_6 + k_7^{\max} \frac{[S]^n}{K^n + [S]^n}} \left[ \frac{1}{4\pi D_s a_s [S]} + \frac{1}{k_2} \right] + \frac{1}{k_3} + \frac{1}{k_4} + \frac{1}{k_5}. \quad (8)$$

#### 2 Supplementary Figures

##### Fig. S1: Slow-Route Landscape and Low-Turnover Rate Profile

In the isolated slow-route landscape, the enzyme cannot enter the catalytic cycle directly from the tensed state. Instead, turnover proceeds through the extended sequence $E_R \rightarrow E_R \cdots S \rightarrow E_T \cdots S \rightarrow ES \rightarrow EP \rightarrow E_T \rightarrow E_R$. Thus, each cycle includes substrate capture by $E_R$, conformational passage across $\Delta G_1^\ddagger$, catalytic progression through $ES$ and $EP$, product release across $\Delta G_3^\ddagger$, and relaxation back to $E_R$ across $\Delta G_4^\ddagger$. Because the faster $E_T + S$ entry route is absent, the system remains confined to the lower turnover branch.

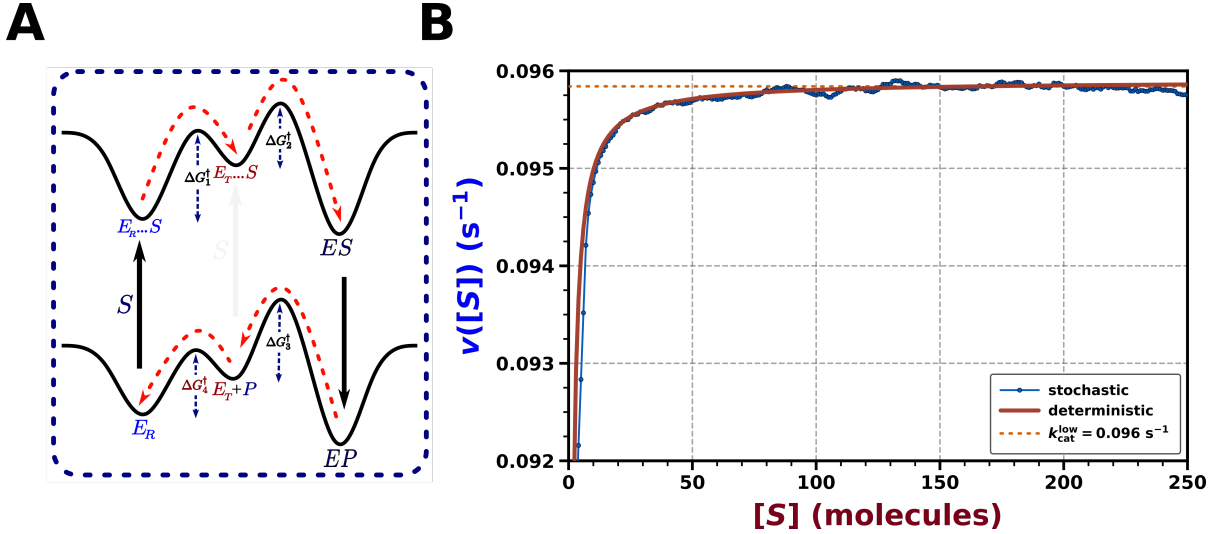

**Fig. S1:** Isolated slow catalytic pathway. (A) Modified free-energy landscape obtained by suppressing the direct substrate-entry route from the tensed enzyme conformation. (B) Velocity-substrate profile showing a single low-turnover saturation regime with $k_{\text{cat}}^{\text{low}} = 0.096 \text{ s}^{-1}$, with close agreement between stochastic and deterministic results.

#### Fig. S2: Fast-Route Landscape and High-Turnover Rate Profile

In this limiting landscape, the relaxed-state branch is removed from the kinetic network, so substrate entry occurs only through the tensed enzyme conformation. The enzyme therefore follows the shortened catalytic route $E_T + S \rightarrow E_T \cdots S \rightarrow ES \rightarrow EP \rightarrow E_T + P$, bypassing the slow relaxation step through $E_R$. Since the recovery bottleneck is absent, catalytic flux remains confined to the fast route, and the rate–substrate response approaches a single high-turnover saturation limit rather than displaying two distinct regimes.

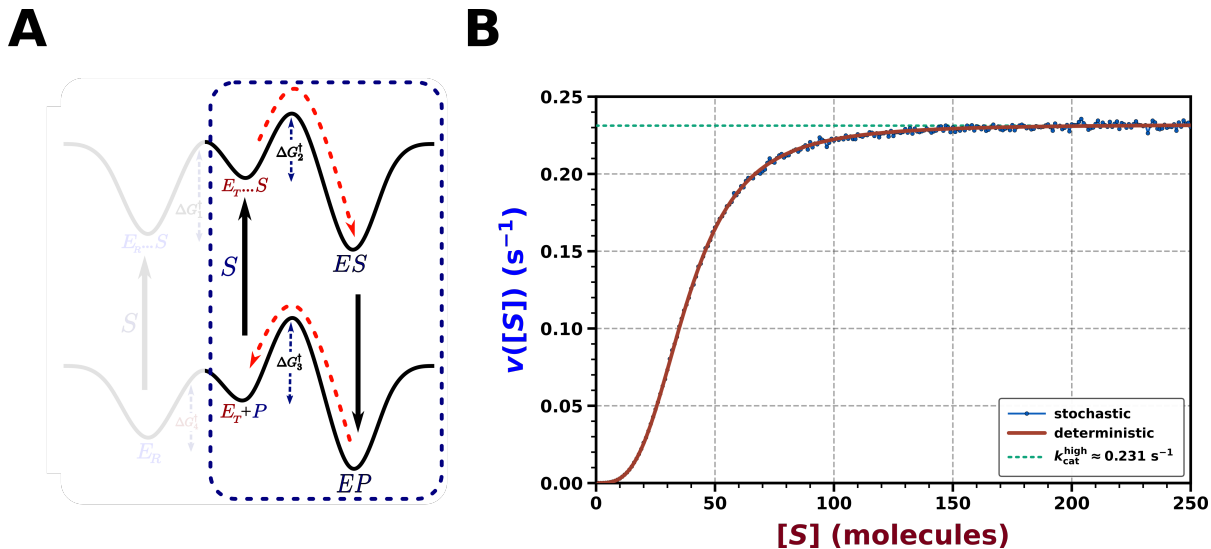

**Fig. S2:** Isolated fast catalytic route. (A) Modified free-energy landscape obtained by removing the relaxed-state branch and retaining direct substrate entry through the tensed enzyme conformation. (B) Velocity–substrate profile showing a single high-turnover saturation limit, $k_{\text{cat}}^{\text{high}} \approx 0.231 \text{ s}^{-1}$, with close agreement between stochastic and deterministic results.

##### Fig. S3: Stochastic Rate Profiles and Shift of the Kinetic Isosbestic Point under Barrier Variation

To further examine the barrier dependence of the kinetic isosbestic point, the relaxation barrier $\Delta G_4^\ddagger$ was varied at fixed values of the substrate-bound conformational barrier $\Delta G_1^\ddagger$. Four fixed values of $\Delta G_1^\ddagger$ were considered: 0.5, 1.0, 1.5, and $2.0 k_B T$. For each of these cases, $\Delta G_4^\ddagger$ was varied over 0.5, 1.0, 1.5, and $2.0 k_B T$, generating a family of velocity–substrate profiles. In every panel, the stochastic curves intersect at a common point, confirming that the isosbestic response is retained when $\Delta G_4^\ddagger$ is modulated. However, increasing $\Delta G_1^\ddagger$ shifts the intersection systematically toward lower substrate concentration and lower catalytic velocity. The isosbestic coordinate moves from approximately $S = 74$, $v = 0.168 \text{ s}^{-1}$ at $\Delta G_1^\ddagger = 0.5 k_B T$ to $S = 46$, $v = 0.086 \text{ s}^{-1}$ at $\Delta G_1^\ddagger = 2.0 k_B T$. This shift reflects the increasing kinetic resistance for the $E_R \cdots S \rightarrow E_T \cdots S$ transition, which enhances the influence of the slower catalytic route.

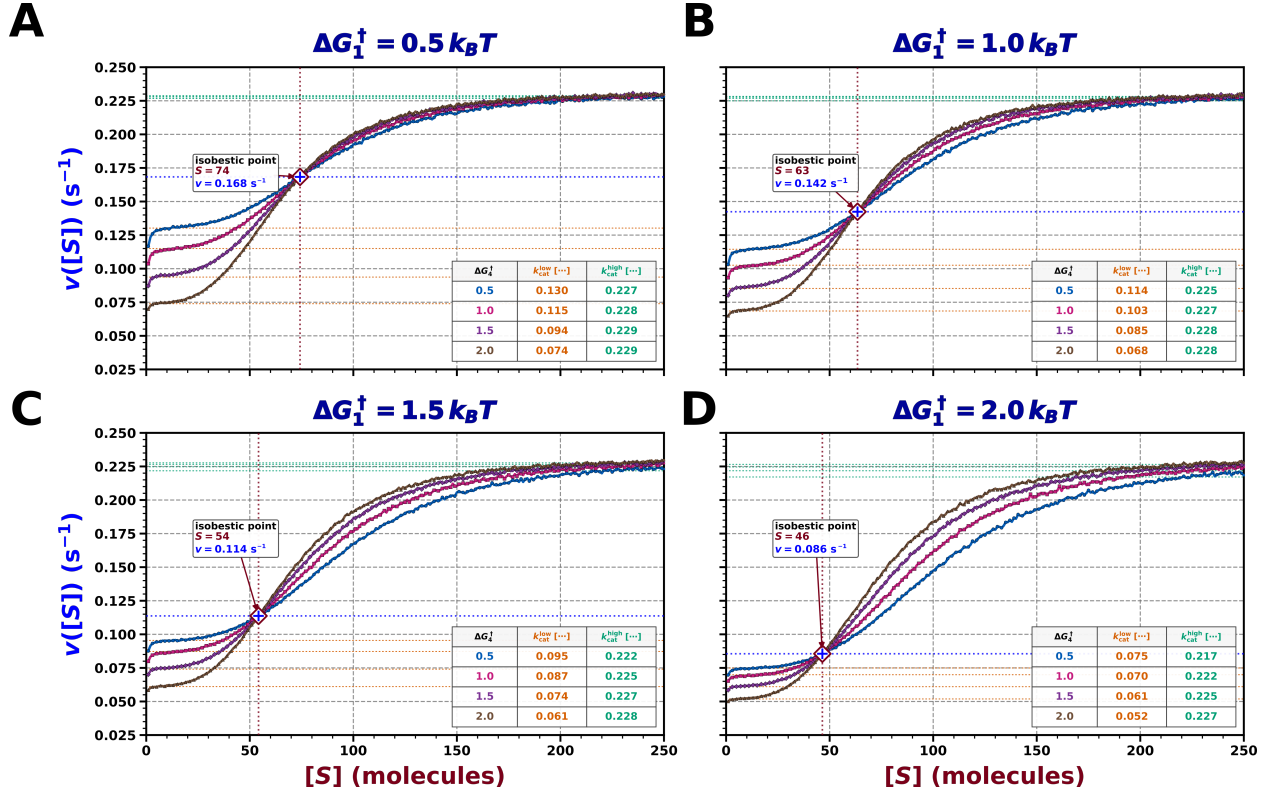

**Fig. S3:** Stochastic isosbestic response under barrier variation. Velocity–substrate profiles are shown for different $\Delta G_4^\ddagger$ values at fixed $\Delta G_1^\ddagger = 0.5, 1.0, 1.5$, and $2.0 k_B T$. The common intersection shifts to lower $S$ and lower $v([S])$ as $\Delta G_1^\ddagger$ increases.

#### Fig. S4: Deterministic Rate Profiles for the Barrier-Dependent Isosbestic Point

The deterministic rate expression in Eq. 7 was used to examine the same barrier-dependent isosbestic behavior shown in Fig. S4. Here, the substrate-bound conformational barrier was fixed at $\Delta G_1^\ddagger = 0.5, 1.0, 1.5,$ and $2.0 k_B T$, while the relaxation barrier $\Delta G_4^\ddagger$ was varied over $0.5, 1.0, 1.5,$ and $2.0 k_B T$. For each fixed $\Delta G_1^\ddagger$, the deterministic velocity–substrate curves intersect at a common point, confirming that the isosbestic response follows directly from the kinetic rate law rather than from stochastic sampling. As $\Delta G_1^\ddagger$ increases, the crossing point shifts from approximately $S = 74, v = 0.168 \text{ s}^{-1}$ to $S = 46, v = 0.085 \text{ s}^{-1}$, showing that the substrate-bound barrier systematically lowers both the crossover substrate level and the corresponding turnover velocity.

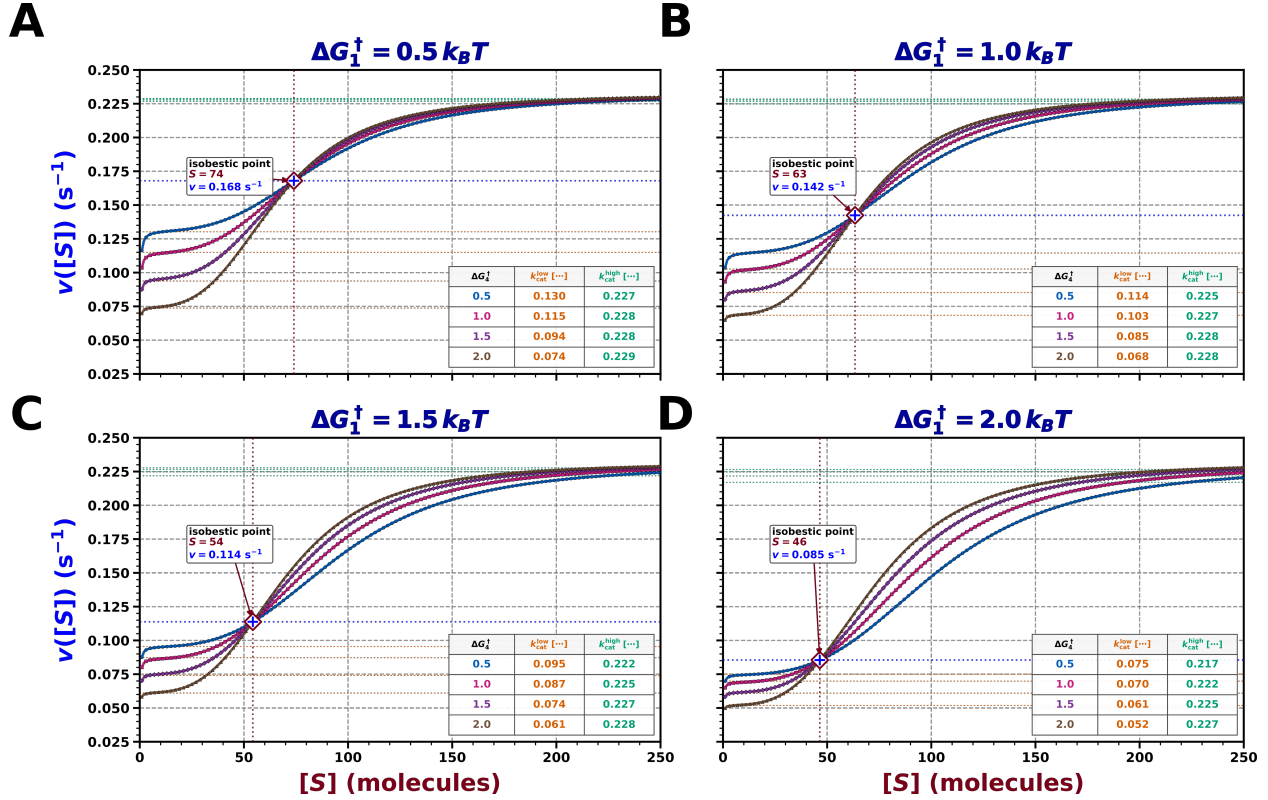

**Fig. S4:** Deterministic isosbestic response under barrier variation. Velocity–substrate profiles are shown for different $\Delta G_4^\ddagger$ values at fixed $\Delta G_1^\ddagger = 0.5, 1.0, 1.5,$ and $2.0 k_B T$. The common crossing point shifts to lower $S$ and lower $v([S])$ as $\Delta G_1^\ddagger$ increases.

**Fig. S5: Scaling of the Isosbestic Coordinates with $\Delta G_1^\ddagger$**

The displacement of the kinetic isosbestic point was quantified by extracting the crossover velocity, $v_c = v(S_c)$, and the corresponding substrate value, $S_c$, for each value of the substrate-bound conformational barrier $\Delta G_1^\ddagger$. The analysis was performed independently for the stochastic and deterministic rate profiles. In both cases, $v_c$ decreases approximately linearly with increasing $\Delta G_1^\ddagger$, whereas $S_c$ follows an exponential decay. The close agreement between the stochastic and deterministic fits confirms that the barrier-dependent shift of the isosbestic point is an intrinsic feature of the kinetic network.

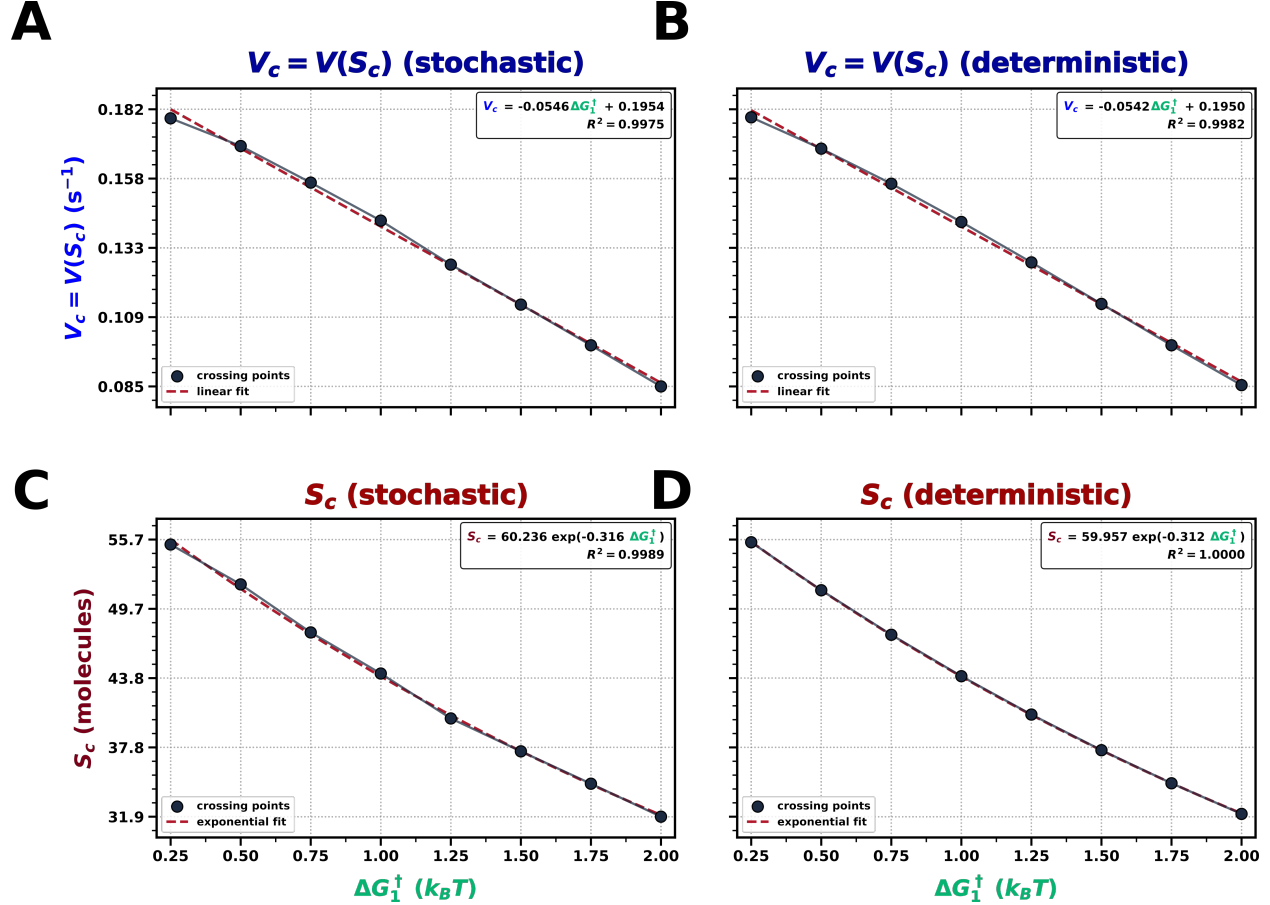

**Fig. S5: Barrier-dependent scaling of the isosbestic coordinates.** (A,B) Linear dependence of the crossover velocity $v_c$ on $\Delta G_1^\ddagger$ for stochastic and deterministic rate profiles. (C,D) Exponential decay of the crossover substrate value $S_c$ with $\Delta G_1^\ddagger$ for stochastic and deterministic descriptions.

##### Fig. S6: Effect of $\Delta G_1^\ddagger$ and $\Delta G_4^\ddagger$ Modulation on Burst–Halt Turnover

To distinguish the effects of the two conformational barriers on single-trajectory product formation, cumulative product trajectories were analyzed at fixed substrate level $S = 500$. In the first case, $\Delta G_4^\ddagger$ was fixed at $1.0 k_B T$, while $\Delta G_1^\ddagger$ was varied from $0.5$ to $5.0 k_B T$. Increasing $\Delta G_1^\ddagger$ produces stronger departures from uniform product accumulation, indicating longer residence in the slow catalytic route. In the second case, $\Delta G_1^\ddagger$ was fixed at $1.5 k_B T$, while $\Delta G_4^\ddagger$ was varied over the same range. These trajectories remain nearly linear and closely grouped, showing that modulation of $\Delta G_4^\ddagger$ does not generate the same burst–halt intermittency.

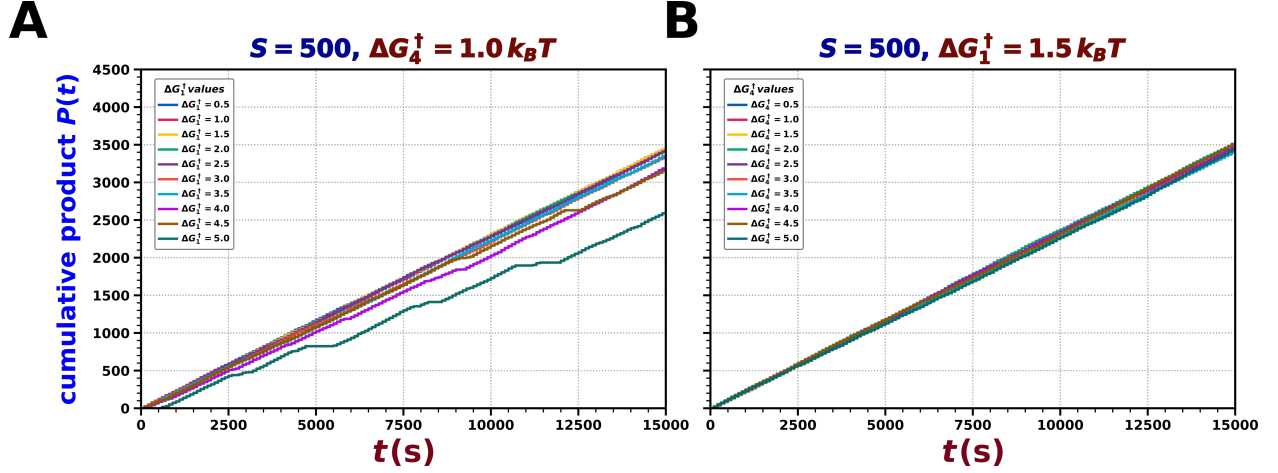

**Fig. S6:** Barrier-specific effects on cumulative product formation at $S = 500$. (A) Product trajectories obtained by varying $\Delta G_1^\ddagger$ at fixed $\Delta G_4^\ddagger = 1.0 k_B T$, showing enhanced intermittency at larger $\Delta G_1^\ddagger$. (B) Product trajectories obtained by varying $\Delta G_4^\ddagger$ at fixed $\Delta G_1^\ddagger = 1.5 k_B T$, showing nearly uniform product accumulation.

#### Fig. S7: Convergence to Michaelis–Menten Kinetics at Asymptotic $\Delta G_4^\ddagger$ Barriers

The deterministic rate expression in Eq. 7 was further examined over an extended substrate range to clarify the limiting behaviors observed in the stochastic profiles. Supplementary Fig. S7A shows the large-barrier condition, $\Delta G_4^\ddagger = 4.0 k_B T$, for which suppression of the $E_T \rightarrow E_R$ relaxation step strongly favors the faster tensed-state pathway and produces an apparent high-rate, single-saturation response over the substrate range considered in the stochastic analysis. Extending the deterministic profile toward much lower substrate levels, however, resolves an additional low-rate saturation regime before the system crosses over to the upper plateau. Conversely, Supplementary Fig. S7B corresponds to the small-barrier condition, $\Delta G_4^\ddagger = 0.04 k_B T$, where rapid relaxation toward $E_R$ favors the slower pathway and yields an apparent low-rate saturation profile over the same range; at sufficiently high substrate levels, the response ultimately departs from this plateau and approaches the higher turnover regime. Supplementary Fig. S7C further shows the corresponding sigmoidal response, in which the transition between the low- and high-rate regimes becomes sharply localized around a characteristic substrate level, reproducing the allosteric-like switching behavior observed in the stochastic profile. Together, these results show that the apparent single-saturation and sigmoidal limits retain the underlying bimodal kinetic structure of the model.

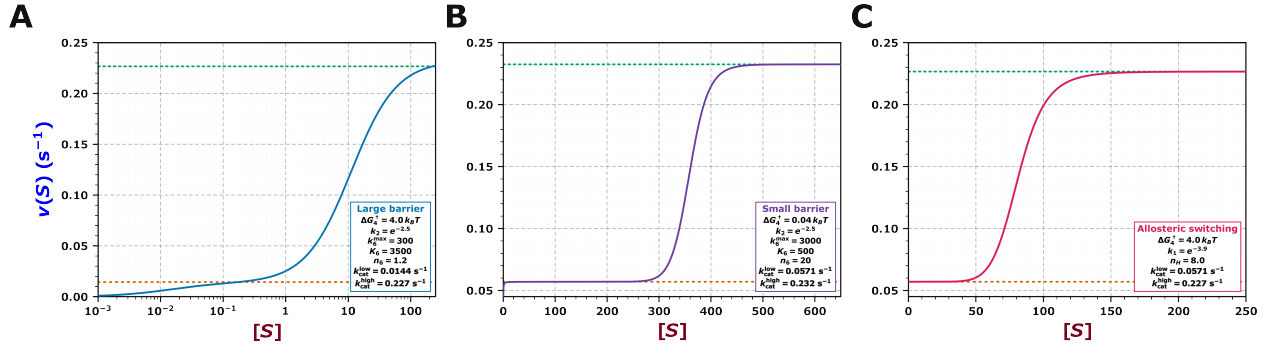

**Fig. S7:** Deterministic rate–substrate profiles across extended substrate ranges. **(A)** Under the large-barrier condition, $\Delta G_4^\ddagger = 4.0 k_B T$, extension toward very low substrate levels resolves the lower-rate regime preceding saturation at the higher turnover limit. **(B)** Under the small-barrier condition, $\Delta G_4^\ddagger = 0.04 k_B T$, extension toward high substrate levels reveals the transition from the lower plateau to the higher turnover regime. **(C)** Parameter modulation produces a pronounced sigmoidal transition between the two saturation regimes, resembling an allosteric-like kinetic response.
